# From Light to Lipids: Constraint-Based Metabolic Modeling of *Nannochloropsis oceanica* Under Light Acclimation Conditions

**DOI:** 10.64898/2025.12.19.695437

**Authors:** Sabine van Oossanen, Narcis Ferrer-Ledo, Sarah D’Adamo, Vitor A.P. Martins Dos Santos, Maria J. Barbosa, Maria Suarez-Diez

## Abstract

Oleaginous microalgae, such as *Nannochloropsis oceanica*, hold strong promise for sustainable lipid bioproduction, but fully realizing this potential requires systems-level insight into their complex metabolism. One of the central challenges in optimizing lipid productivity is the resolution of the highly heterogeneous and incompletely annotated metabolic networks, which respond dynamically to strain-specific traits and cultivation conditions. Here we present iSO1949_N.oceanica, the first genome-scale constraint-based metabolic model (GEM) for this species. Constructed through an orthology-based approach using curated models of related microalgae, the GEM integrates core photosynthetic metabolism and lipid biosynthesis pathways with extensive subcellular localization predictions. To capture environmental dynamics, we introduce two light-acclimation modes derived from continuous cyclostat cultivations, incorporating biomass composition, oxygen exchange, and maintenance rates based on photosynthesis–irradiance curves. Simulations reproduce carbon assimilation under variable light conditions and differentiate acclimated phenotypes. iSO1949_N.oceanica provides a comprehensive framework for exploring photosynthetic metabolism and guiding engineering strategies under photobioreactor-relevant conditions. This resource advances the use of *N. oceanica* as a chassis for sustainable lipid production and establishes a foundation for systems-level analysis of stramenopile microalgae.

## 1 Introduction

Sustainable oil sources are essential for food, feed, and fuel production in a biobased economy. Compared to conventional oil sources, oleaginous microalgae, such as *Nannochloropsis oceanica*, show promise since these do not compete with traditional food crops for space and substrates, can achieve lipid contents exceeding 60% g/gDW under nutrient limiting conditions (Ma et al., 2016), and can produce significant portions in the form of the high-value omega-3 fatty acid eicosapentaenoic acid (EPA).

To fulfill the demands of current primary lipid sources like fish or palm oil, algal lipid productivities must increase (Ashour et al., 2019; Barbosa et al., 2023; Chauton et al., 2014; Nagappan et al., 2021). Metabolic engineering is an upcoming approach for increasing native lipid productivities, facilitated by recent genomic and genetic tool developments (Janssen et al., 2020; Naduthodi et al., 2021, 2019; Radakovits et al., 2012; Südfeld et al., 2022; Vieler et al., 2012; Wei et al., 2013). Current approaches to selecting genetic targets for increasing lipid productivity are limited, focusing on the metabolism of high neutral lipid accumulating phenotypes. An increasing body of transcriptomics, proteomics, and lipidomics datasets has been obtained under diverse cultivation conditions that can aid in target selection (Dong et al., 2013; Li et al., 2014; Wang and Jia, 2020). High-lipid phenotypes are generally derived from stress-inducing conditions such as nitrogen deprivation or light-stress (Han et al., 2017; Wang and Jia, 2020). These phenotypes show hampered growth and photosynthetic efficiency. As such, targets derived from these phenotypes might not result in viable and robust lipid productivities, limiting industrial applicability.

Computational approaches, such as genome-scale constraint-based metabolic models (GEMs), support rational selection of metabolic targets to improve lipid productivities (Tibocha-Bonilla et al., 2018). GEMs map all annotated metabolic conversions within an organism, simulating active pathways under varying conditions through flux balance analysis (FBA). Owing to the link between annotated genes and metabolic reactions, the GEM can assess *in-silico* how each gene affects the production of metabolites of interest, allowing for faster, more efficient, and more impactful selection of metabolic targets. GEMs have shown potential in increasing lipid production in oleaginous yeast (Kim et al., 2019; Pham et al., 2021) and more than 40 GEMs have been developed for over 15 microalgal species. Current GEMs capture key algal functionalities including light-dependent energy production and nitrogen starvation responses, though experimental validation has been limited. Most microalgal GEMs have been developed for model organisms such as *Chlamydomonas reinhardtii* (Imam et al., 2015; Kliphuis et al., 2012), *Phaeodactylum tricornutum* (Broddrick et al., 2022; Levering et al., 2016), and the model cyanobacteria *Synechocystis* (Tibocha-Bonilla et al., 2018), for which high-quality genome annotations and specific genetic tools are available. Moreover, GEMs have also been developed for *Microchloropsis salina* (Shah et al., 2017) and *Microchloropsis gaditana* (Dupont-Thibert et al., 2023; Loira et al., 2017), formerly classified as part of the *Nannochloropsis* genus (Fawley et al., 2015). Despite its industrial relevance, no GEM exists for *Nannochloropsis oceanica*.

The predictive capability of microalgal GEMs is currently limited by several gaps, particularly in metabolic understanding, in protein annotation and localization, and by the dynamic nature of light capture mechanisms. Light intensity and its fluctuations significantly affect biochemical composition, carbon partitioning, and biomass yields of microalga, posing challenges for GEMs to accurately represent these effects metabolic effects (Alboresi et al., 2016; Perin et al., 2023; Wang and Jia, 2020). Moreover, available datasets on biomass-specific light supply, acclimation, and biomass compositions are limited. Most studies report incident light intensities and photoperiod, without accounting for the perceived light by microalgae throughout the culture, which depends on culture density, light absorption, and light path length. A recent study explored the effect of both the incident light intensity in combination with light gradients on biomass physiology and lipid profile of *Nannochloropsis oceanica* in long-term acclimated cultures (Ferrer-Ledo et al., 2025). This study provides a valuable phenotypic dataset that can be used to refine a GEM with light acclimated constraints.

Another limiting factor for algal GEMs is limited model reusability, due to inconsistent metabolite IDs, ambiguous database references, and untransparent curation processes (Pham et al., 2019). Applying the principles of Findability, Accessibility, Interoperability, and Reusability (FAIR) during the model construction process can mitigate these issues, enhancing the reusability and reliability of algal GEMs (Wilkinson et al., 2016).

In this work, we present a novel GEM for *N. oceanica* called iSO1949_N.oceanica, curated in detail on the lipid metabolism under long-term high and low light acclimated conditions. The iSO1949_N.oceanica GEM is developed in a FAIR, traceable orthology-based approach, using four individually developed algal GEMs to prevent bias transfer. In addition, iSO1949_N.oceanica includes state-of-the-art protein localization predictions. This model offers reliable gene localization and the exploration of putatively annotated genes and reactions. Through comparison with extensive biomass compositions and photosynthetic response data, this model has been shown to predict the impact of light intensity and light acclimation on carbon fixation and energy requirement.

## 2 Methods

### 2.1 Pipeline for protein localization

Protein localizations were assigned for all annotated nuclear encoded proteins through a pipeline from (Levering et al., 2016), supplemented with prediction tools ASAFind, ASAFind 2.0, and DeepLoc2.0. We obtained protein sequences of *Nannochloropsis oceanica* IMET1 from the NanDeSyn database (https://nandesyn.single-cell.cn/download, IMET1v2.protein.faa, Gong et al., 2020), as input for DeepLoc 2.0 (Thumuluri et al., 2022), HECTAR 1.3 (Gschloessl et al., 2008), SignalP4.1 through 6.0 (Almagro Armenteros et al., 2019; Nielsen, 2017; Teufel et al., 2022), TargetP 2.0 (Armenteros et al., 2019), ASAFind (Gruber et al., 2015), ASAFind2.0 (Gruber et al., 2025), PredictNLS (Cokol et al., 2000), and MitoFates (Fukasawa et al., 2015). Each tool was used for localization prediction using the settings listed in Data S2. In addition, each protein sequence was scanned for plastidic, endoplasmic and peroxisomal sequence motives. If a specific location was predicted by multiple tools or sequence motives as indicated in Figure 1B, the localization score was increased. The organelle with the highest localization score was selected. In case of equal compartment scores, a location was assigned based on compartment hierarchy as shown in Figure 1A and described in the Appendix S1A. The prediction scores per compartment have been provided in Data S2, which allows users to investigate individual scores and to apply alternative strategies for compartment allocation to their preference.

**Figure 1.**
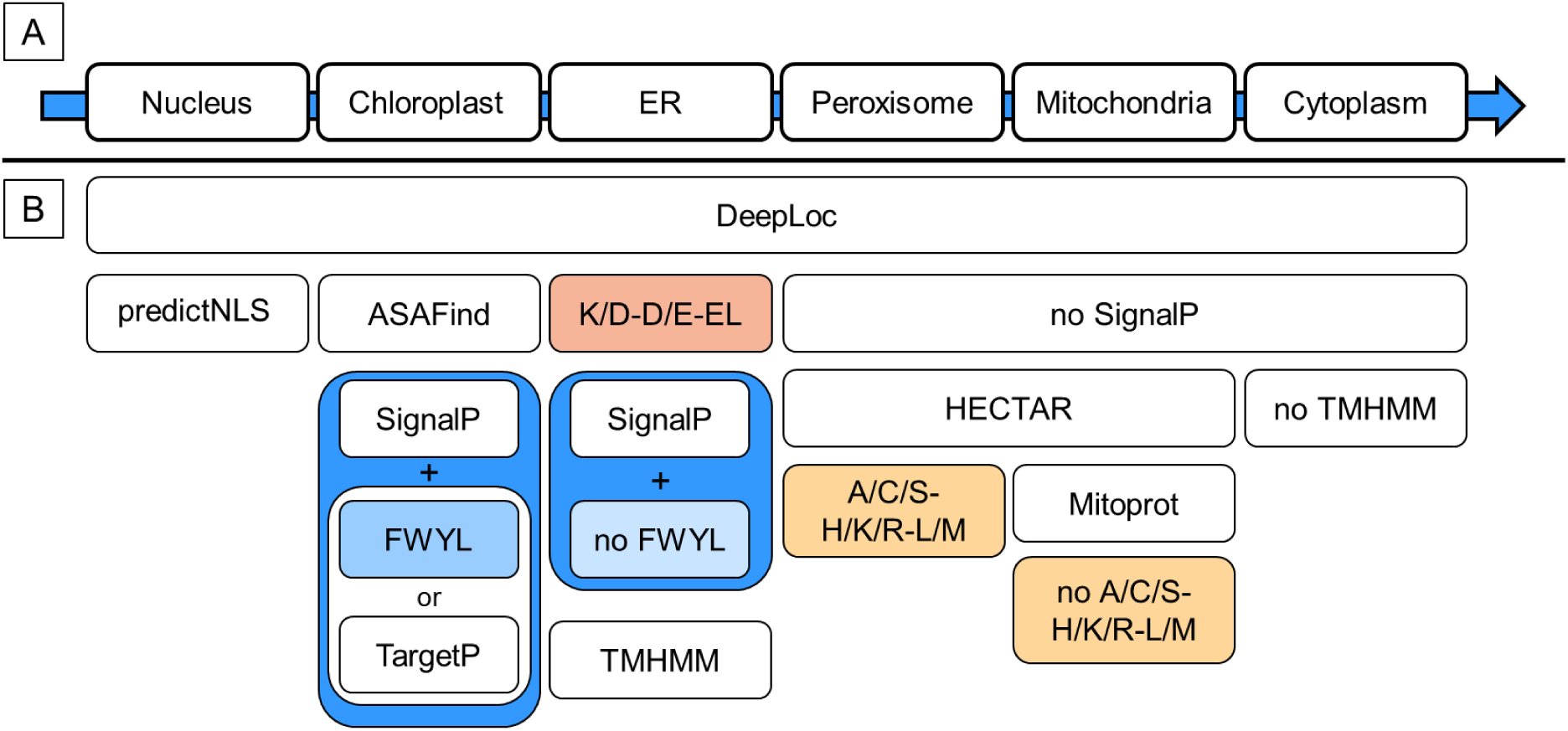
(A,B) Pipeline for predicting protein localization of annotated Nannochloropsis oceanica IMET1 genes, organized per organelle. White boxes in (B) represent which tools were used to predict localization in the organelles indicated in (A), and colored text boxes represent whether a signal peptide should be present or absent. The more tools predict localization of the protein in a specific compartment, the higher the localization score. The protein was assigned to the organelle with the highest localization score, sorted as indicated by the arrow.

### 2.2 Phenotype data retrieval

Phenotypic data from *Nannochloropsis oceanica spp.* was obtained from samples grown in cyclostat conditions under varying light intensities and light gradients as reported previously (Ferrer-Ledo et al., 2025). The data is extensively described by Ferrer-Ledo, van Oossanen et al.(Ferrer-Ledo et al., 2025), to where we refer the reader for extensive information. In brief, *Nannochloropsis oceanica spp*. was grown under a 16:8h light:dark regime and in semi-continuous cyclostat conditions. Cyclostats have constant dilution during the light period and no dilution during the dark period. We selected data from replicate low light (LL) cyclostats with incident light intensity (I_ph,0_) of 200 µmol m^-2^ s^-1^ and an average specific light absorption rate (q_ph,abs_) of 20.8 mmol_ph_ gDW^-1^ h^-1^, in addition to high light (HL) cyclostats with an I_ph,0_ of 670 µmol m^-2^ s^-1^ and q_ph,abs_ of 69.3 mmol_ph_ gDW^-1^ h^-1^. Samples for PI-curves were taken from the reactor precisely two hours into the light period. Samples for biomass biochemical analysis were taken from the overflow, which was collected between the second hour and the sixth hour of the light period and was stored on ice upon leaving the reactor.

### 2.3 Biochemical analysis

Biomass composition analysis was performed on *N. oceanica spp.* samples grown in cyclostat conditions under varying light intensities and light gradients (Ferrer-Ledo et al., 2025). Per cyclostat setting, three freeze-dried biomass samples from individual steady state days were obtained. We analyzed total carbohydrate content according to the Dubois colorimetric method (Dubois et al., 1956) and total protein content according to the Lowry method (Lowry et al., 1951). Extraction and quantification of both were performed as described in (Gao et al., 2021), with minor edits. Carbohydrate extraction was modified by adding 5mL of 2.5M NaOH solution after boiling the acidic biomass solution for 3 hours, followed by thorough vortexing. Protein extraction was modified by analyzing 4.5-11.5 g biomass, resulting in a protein concentration range of both the carbohydrate and protein extractions were performed in duplicate per biomass sample, followed by duplicate quantification with sulfuric acid – phenol solutions, resulting in 4 data points per biomass sample, and 12 points per cyclostat setting.

### 2.4 Orthology-based GEM construction

Orthology-based GEM reconstruction was performed using the AuReMe automatic tool (Aite et al., 2018). Genomic input for *N. oceanica* IMET1 was obtained from the NanDeSyn database (https://nandesyn.single-cell.cn/download, file IMET1v2.protein.faa), supplemented by plastidial and mitochondrial encoded protein sequences from Genbank (accession numbers KC568456, KC598086, Wei et al., 2013). As scaffolds for orthologous reconstruction, GEMs of *Microchloropsis salina* (iNS934, (Loira et al., 2017)), *Microchloropsis gaditana* (iRJ1321, Shah et al., 2017), *Chlamydomonas reinhardtii* (iCre1355, Imam et al., 2015), and *Phaeodactylum tricornutum* (iLB1027_lipid, Levering et al., 2016) were used. Specifics on protein sequences and SBML file retrieval and pretreatment are detailed in Appendix S1A. The AuReMe settings and protocol used to generate the draft GEM from the scaffold GEMs can be found on GitLub (https://gitlab.com/wurssb/Modelling/i-so-1949-n-oceanica-gem).

Metabolite IDs of the resulting draft were mapped to BiGG (Norsigian et al., 2020), KEGG (Kanehisa et al., 2017; Kanehisa and Goto, 2000), ModelSEED (Henry et al., 2010) and MetaCyc (Karp et al., 2019) and supplemented with InchiKeys (Goodman et al., 2021; Heller et al., 2015) and chemical formulas using the BiGG metabolite database, supplemented by the MetaCyc Metabolite Translation Service, ModelSEED compounds database, and MetaNetX (Moretti et al., 2021). Details on the mapping process are provided in Appendix S1A.3. When metabolite IDs could not be retrieved from any database, or when automatic mappings conflicted, IDs were looked up and mapped manually. Finally, conflicting mappings, missing annotation and structural annotations were solved through manual curation, specifically for lipid species. In case of doubt, molecular structures were compared on KEGG and ModelSEED. Metabolite labels for lipid class and acyl-chain length were added for each lipid species to improve readability of the GEM. Unsaturated lipid metabolites with similar lengths were curated manually and assigned isomer labels, using MetaCyc and ModelSEED as reference. The final SBML file contains the BiGG ID as metabolite name when present, otherwise the ID from KEGG, ModelSEED or MetaCyc would be assigned in that order. Both alternative ID mappings and lipid species labels were added to the metabolites as ‘annotation’ and ‘notes’, respectively.

During the orthology-based reconstruction process several BiGG reactions were replaced by synonymous BiGG reactions carrying alternative reaction equations and alternative compartments. Erroneous BiGG reaction mappings from AuReMe were removed by comparing reaction IDs and equations from the source GEMs to those in the draft GEM. Next, reaction formulas were updated with metabolite mappings and metabolites on both sides of the formula were ordered alphabetically. The ordering of the metabolites was used to flag duplicate reactions for removal. In case both reversible and irreversible versions of a reaction were present, the reversible reaction was kept. In addition, duplicated reactions were evaluated to identify alternative versions with differently protonated variants of substrates or products. Reactions performing a conversion with the same metabolites but in a different stoichiometry or with only a proton difference were curated manually. Gene-Protein-Reactions (GPR) of duplicate reactions were merged with an ‘OR’ relationship. Compartments not relevant to the lipid metabolism or absent in *Nannochloropsis* were excluded, for example the Golgi apparatus, eyespot and flagellum. Alternative IDs found through mapping were added to the metabolite annotation. In addition, notes on lipid acyl-type and lipid class were added for each lipid metabolite. Reactions were annotated with KEGG IDs when present, and reaction notes were added for each individual reaction on their scaffold(s) of origin and on protein localization predictions.

Gap-filling for missing or passive reactions was initiated by including all non-GPR associated transport and export reactions from the scaffold GEMs, to ensure metabolite flow between the compartments. Since *N. oceanica* is known as a strict photoautotroph, import reactions for carbon-containing metabolites and hydrogen (H_2_) were removed except for CO_2_ uptake. Core metabolic pathways and biosynthetic pathways for biomass components were manually curated by comparing the draft GEM with available literature, the iLB1027_lipid GEM and with corresponding KEGG pathway maps. Lipid biosynthesis pathways were tailored to the detected species under varying ingoing light intensities and light gradients. Reactions and metabolites were added to account for species that were detected in the cyclostat studies but were absent from the scaffold GEMs. In addition, lipid species with mixed acyl groups were split into two metabolites carrying a single acyl group since the cyclostat lipid analysis was limited to acyl group distributions over each lipid class. Lipid species that were not detected were removed from the GEM.

### 2.5 Biomass equation

To simulate growth rates for differently light acclimated *N. oceanica*, we added to the GEM biomass-synthesis reactions for high and low light gradient phenotypes grown in semi-steady state cyclostats. Biomass reactions were constructed based on protein, carbohydrate, pigment and lipid contents and compositions from biomass samples taken one hour into the light period. Detailed pigment and lipid contents and compositions are reported by Ferrer-Ledo, van Oossanen et al.(Ferrer-Ledo et al., 2025). In the GEM we included lipid species that amounted to at least 1 nmol/gDW. The total carbohydrate content was combined with a monomeric sugar composition taken from (Jia et al., 2015), combining the free and polymeric sugars at day 4 of nitrogen replete growth. Data was scraped from the article using https://www.graphreader.com/. The total protein content was combined with an amino acid composition taken from *N. oceanica* grown at a high pCO_2_ supply of 1000 ppm (Liang et al., 2020), supplemented with tryptophan contents from *N. oceanica* grown in chemostat mode with 2.2 mM KNO_3_ supply (Xiao et al., 2013). Carbohydrate and protein compositions were used for both the LL and HL biomass reactions. Nucleotide contents and compositions of DNA and RNA fractions were estimated based on the genome size and GC content (Gong et al., 2020), in addition to a DNA:RNA ratio as detected by Rebolloso-Fuentes et al. (Rebolloso-Fuentes et al., 2001) (Data S4E).

Biomass fraction reactions for LL and HL were set up by combining individual metabolites into biomass fraction metabolites, e.g. polar lipids (BM_PL), neutral lipids (BM_NL), carbohydrates (BM_carb), protein (BM_protein) and pigments (BM_pigments). Stoichiometry of these biomass fraction reactions is expressed in molar compositions, using mmol/gDW as unit for the substrate metabolites and g/gDW for the produced biomass fraction metabolites. The biomass fractions were then combined into an overall biomass reaction in a 1:1 ratio. In addition, we accounted for growth associated maintenance in the form of 30 ATP, similar to iNS934 and iCre1355. Detailed calculations of the biomass equation can be found in Data S4.

### 2.6 Photosynthesis measurements

Photosynthesis-irradiance (PI) curves were measured based on dissolved O_2_ using a Biological Oxygen Monitor (BOM) (Oxytherm^+^, Hansatech) and based on dissolved CO_2_ using the LI-6800 (LI-COR Biosciences) as described by (Barten et al., 2022; Yoshida et al., 2023) except for temperature settings which were 25°C for both systems. The PI-curves measure the net O_2_ and CO_2_ production rate (r_O2_, r_CO2_) as a function of incident light intensities (I_ph,0_). O_2_-based curves were measured for replicate steady state samples of each reactor described by Ferrer-Ledo, van Oossanen et al. (Ferrer-Ledo et al., 2025), whereas CO_2_-based curves were measured using only one replicate reactor of each light condition. After sampling the reactor, samples were stored in the dark and inserted into the photosynthesis monitors within 30 minutes.

Biomass specific light absorption rates (q_ph,abs_, mmol gDW^-1^ h^-1^) were calculated as below:

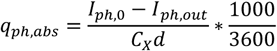

Where I_ph,0_, I_ph,out_, C_X_, and d represent the ingoing and outgoing light intensity (µmol m^-2^ s^-1^), the biomass concentration (gDW m^-3^), and light path (m) within the sample cuvette. The light path was simplified for the BOM, which contains a circular cuvette. Since the light path varies in a circular cuvette, we recalculated the light path by refitting the cuvette surface into a square, and using the square diameter. The I_ph,out_ was determined as below:

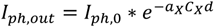

Where a_X_ represents the absorption cross-section (m^2^ kg^-1^). The specific O_2_ conversion rates (q_O2_) and CO_2_ conversion rates (q_CO2_) were determined based on the net O_2_ and CO_2_ production rates (r_O2_, r_CO2_, mmol m^-3^ h^-1^) obtained from the PI curves and C_X_ as follows:

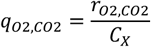

Maximum q_O2_ values per reactor were calculated by averaging the maximum q_O2_ of 2 or 4 replicate steady state PI-curves. For the LL reactors, one reactor replicate was excluded based on observed photo-inhibition before reaching the maximal q_O2_. Respiration rates were calculated as the intercept of the light-limited part of the oxygen-based PI-curve, before photoinhibition was observed (≤ 61 µmol_ph_ m^-2^ s^-1^). Average respiration rates were calculated for n=2 and n=5 PI-curves of the low light replicate reactors, and n=4 for each high light replicate reactor. Respiration rates from the reactors were averaged to obtain the LL and HL respiration rates, which were used to constrain the GEM non-growth associated maintenance (NGAM) reaction ATPPH.

### 2.7 GEM simulations

The GEM was modified and analyzed in Python v3.7.1 using COBRApy v0.13.4 (Ebrahim et al., 2013) and Gurobi as solver (Gurobi Optimization, LLC, 2023). Nutrient uptake was restricted to CO_2_ and NO_3_ as carbon and nitrogen source, respectively. Additional constraints were a forced cyclic electron flow of 0.3 mmol gDW^-1^ h^-1^, and a limited demand of mitochondrial formaldehyde (DM_fald_m) of 0.1 mmol gDW^-1^ h^-1^. NO_3_ uptake and O_2_ production were limited to 5.0 and 6.5 mmol gDW^-1^ h^-1^ respectively. Neutral lipid (NL) sink reactions specific to the LL or HL lipid compositions were added to allow simulation of carbon partitioning between biomass and NL under the varying light conditions.

PI-curves were simulated by varying the photon uptake rate between 0-200 mmol gDW^-1^ h^-1^, maximizing CO_2_ uptake, and calculating the corresponding O_2_ exchange ranges, growth rates and NL accumulation rates through flux variability analysis (FVA) with loopless settings. PI-curves were adjusted to the acclimated LL and HL phenotypes by constraining maximum O_2_ production, NGAM, biomass composition and NL composition to the experimentally determined values per phenotype. Under light supply rates below the light compensation point, where no net O_2_ is produced, intracellular glucose was supplied at 0.12 mmol gDW^-1^ h^-1^ to allow for respiration.

## 3 Results and Discussion

### 3.1 GEM overview

In this study, we constructed the first comprehensive genome-scale constraint-based metabolic model (GEM) for *Nannochloropsis oceanica* IMET1 (Data S2 and S3). The construction process was divided into 3 phases (Figure 2). First, a draft GEM was constructed through orthology-based reconstruction with AuReMe, using published scaffold GEMs from *M. salina, M. gaditana, C. reinhardtii* and *P. tricornutum*. The resulting draft GEM contained 1977 genes, 4781 metabolites and 5970 reactions. Second, the draft was curated by i) mapping metabolite IDs both automatically and manually to well-known metabolite databases BiGG, KEGG, modelSEED and MetaCyc, ii) removing double and faulty reactions and incorrect or incomplete compartments such as the eye-spot, flagellum and Golgi, iii) gap-filling transport reactions and missing reactions of core metabolic pathways focusing on biosynthesis of lipids and main biomass components, and iv) adding localization predictions labels to each reaction with GPR, and lipid annotation labels to each lipid metabolite. Finally, biomass reactions were added from compositions of high light (HL) and low light (LL) adapted cultures. The resulting GEM contained 1949 genes, 3291 metabolites and 3491 reactions (Table 1). An overview of the metabolites and reactions that were removed or added during the curation process, as well as the finally included metabolites and reactions is provided in Data S3. MEMOTE (Lieven et al., 2020) was used to evaluate the quality of the model and a score of 27% was obtained, which is higher compared to the scaffold GEMs that scored between 14-26%. The consistency score of 49% and metabolite annotation score of 63% reflect the high quality and high level of metabolite curation present in the GEM (Appendix S2).

**Figure 2.**
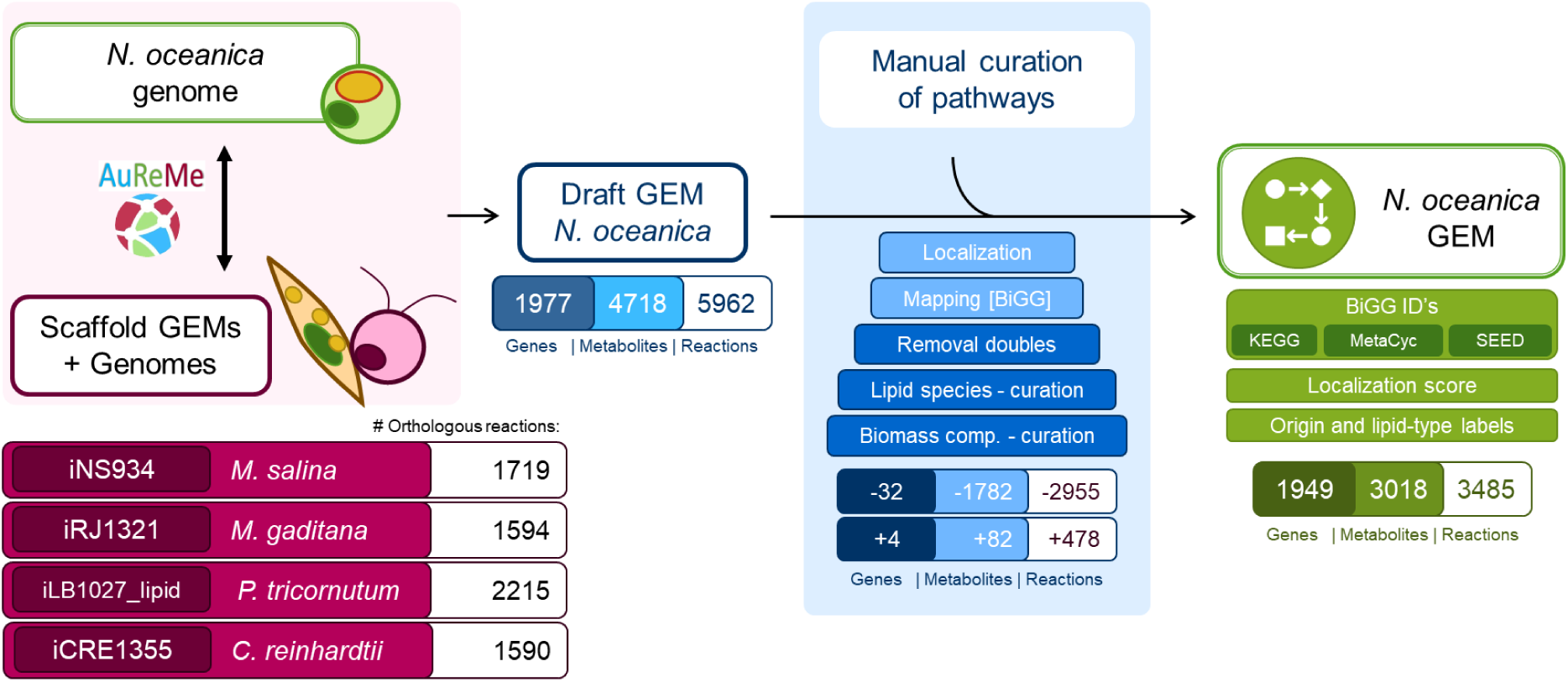
Orthology-based reconstruction process of the iSO1949_N.oceanica model.

**Table 1.** Properties of GEM compared to scaffold GEMs.

| Model | ISO1949_N.oceanica<br>(this work) | iNS934<br>(Loira et al., 2017) | iRJ1321<br>(Shah et al., 2017) | iLB1027_lipid<br>(Levering et al., 2016) | iCre1355<br>(Imam et al., 2015) |
| --- | --- | --- | --- | --- | --- |
| Organism | *N. oceanica* IMET1 | *M. salina* | *M. gaditana* | *P. tricornutum* | *C. reinhardtii* |
| Namespace | BiGG | BiGG | BiGG, Metacyc | BiGG | BiGG |
| Scaffold and reference GEMs | iNS934, iRJ1321, iLB1027_lipid, iCre1355 | iRC1080 (Chaiboonchoe et al., 2014) | NagaCyc, iRC1080, iCZ843 (Zuñiga et al., 2016), iJN678 (Nogales et al., 2012) | iRC1080, iJN678, Knoop GEM (Knoop et al., 2013) | iRC1080, ChlamyCyc (Plant Metabolics Network (PMN), 2014) |
| Protein encoding genes in genome | 12442 | 17519 | 10929 | 22893 | 19526 |
| Genes in GEM (% of annotated) | 1949 (15.9%) | 934 (5.3%) | 1321 (12.1%) | 1027 (4.5%) | 1355 (6.9%) |
| Compartments | 8 | 10 | 4 | 6 | 11 |
|  | extracellular, cytosol, mitochondria, chloroplast, thylakoid, endoplasmic reticulum, peroxisome, nucleus | extracellular, cytosol, mitochondria, chloroplast, thylakoid, endoplasmic reticulum, peroxisome, Golgi apparatus, lysosome, nucleus | extracellular, cytosol, mitochondria, chloroplast | extracellular, cytosol, mitochondria, chloroplast, thylakoid, peroxisome | extracellular, cytosol, mitochondria, chloroplast, thylakoid, peroxisome, Golgi apparatus, lysosome, nucleus, eyespot, flagellum |
| Reactions / Metabolites | 3485 / 3081 | 2345 / 1985 | 1918 / 1862 | 4456 / 2172 | 2394 / 1845 |
| Unique metabolites | 1714 | 1154 | 1136 | 1583 | 1133 |
| Gene-associated reactions | 3178 | 1736 | 1659 | 4150 | 1862 |
| Non-gene-associated reactions (enzymatic and transport) | 309 | 637 | 143 | 255 | 114 |
| Exchanges with the media, demands and sinks, biomass reactions (total) | 14, 1, 0, 2 (17) | 64, 26, 0, 4 (94) | 36, 3, 20, 1 (60) | 30, 11, 3, 7 (51) | 55, 36, 0, 3 (94) |

The final model, iSO1949_N.oceanica, can be found in table format as Data S3 and in SBML L3V1 standardization (Hucka et al., 2003) as Data S2. Furthermore, the different formats together with a MEMOTE (Lieven et al., 2020) and FROG report (Raman et al., 2024) (https://www.ebi.ac.uk/biomodels/curation/fbc) were combined in an OMEX archive file (Bergmann et al., 2014), deposited in BioModels (Malik-Sheriff et al., 2020), and assigned the identifier MODEL2503190001. Moreover, the model can be accessed on GitLub in StandardGEM format on https://gitlab.com/wurssb/Modelling/i-so-1949-n-oceanica-gem (Anton et al., 2023) where community efforts for improving the GEM can be added continuously.

### 3.2 Orthology-based GEM construction

The automatic metabolic model reconstruction tool AuReMe has been recommended for reconstruction of eukaryotic models in a traceable and FAIR (Findable, Accessible, Interoperable, and Reusable) approach, and it has been deployed to reconstruct microalgal models (Aite et al., 2018; Mendoza et al., 2019). We selected four microalgal scaffold GEMs to use as basis for the draft *N. oceanica* GEM from closely related species and model algae, including *Microchloropsis salina* (iNS934) and *Microchloropsis gaditana* (iRJ1321), both formerly assigned to the *Nannochloropsis* clade (Fawley et al., 2015). Additionally, we used GEMs of model green alga *C. reinhardtii* and diatom *P. tricornutum*. *P. tricornutum,* like *Nannochloropsis*, belongs to the taxonomic group of stramenopiles. Multiple GEMs were available for *C. reinhardtii* and *P. tricornutum*. We selected iCre1355 and iLB1027_lipid based on the curation level of the central carbon metabolism and inclusion of detailed lipid biosynthesis pathways.

Using Aureme, we identified orthologues between the genomes of these scaffold species and *N. oceanica* IMET1. AuReMe then constructed a draft GEM based on the curated reactions linked to these orthologues. An overview of the overview of each scaffold model in terms of reactions and metabolites is provided in Table 2. Notably, the *Microchloropsis gaditana* GEM contributed the most reactions (1594), emphasizing its relevance to *N. oceanica*.

**Table 2.** Reaction contributions from each scaffold model to the draft *N. oceanica* model based on orthology. Reactions with GPR indicate that the reaction is linked to one or more genes, through a Gene-Protein-Reaction annotation. Reactions with GPR can be transferred to a draft GEM based on orthology with the GPR associated gene, and the genome of the organism of interest.

| Scaffold GEM | iRJ1321 | iNS934 | iLB1027 | iCre1355 |
| --- | --- | --- | --- | --- |
| Scaffold organism | *M. gaditana* | *M. salina* | *P. tricornutum* | *C. reinhardtii* |
| No. reactions in scaffold model | 1918 | 2345 | 4456 | 2394 |
| No. reactions with GPR | 1659 | 1741 | 4150 | 1924 |
| No. reactions contributed to GEM draft | 1594 | 1719 | 2215 | 1590 |
| Ratio of reactions with orthology | 96% | 99% | 53% | 83% |
| No. metabolites contributed to GEM draft | 1739 | 1756 | 1992 | 1545 |

We extensively curated the draft GEM by removing redundant and conflicting metabolites and reactions, simplifying or extending lipid biosynthesis pathways, and remapping metabolite and reaction IDs. The curation process is described in detail in Appendix S1D. The finalized GEM contained 3485 reactions, of which 1989 are supported by orthology with a single scaffold organism, and 1271 by orthology with multiple scaffold organisms (Figure 3). Overlap and connectivity between the scaffold reactions was strongly increased during manual curation, particularly owing to integration of lipid metabolism reactions, shown in Appendix S1.3. These efforts demonstrate the added value of using orthology-based reconstruction and FAIR principles in developing a high-quality GEM for *N. oceanica*.

**Figure 3.**
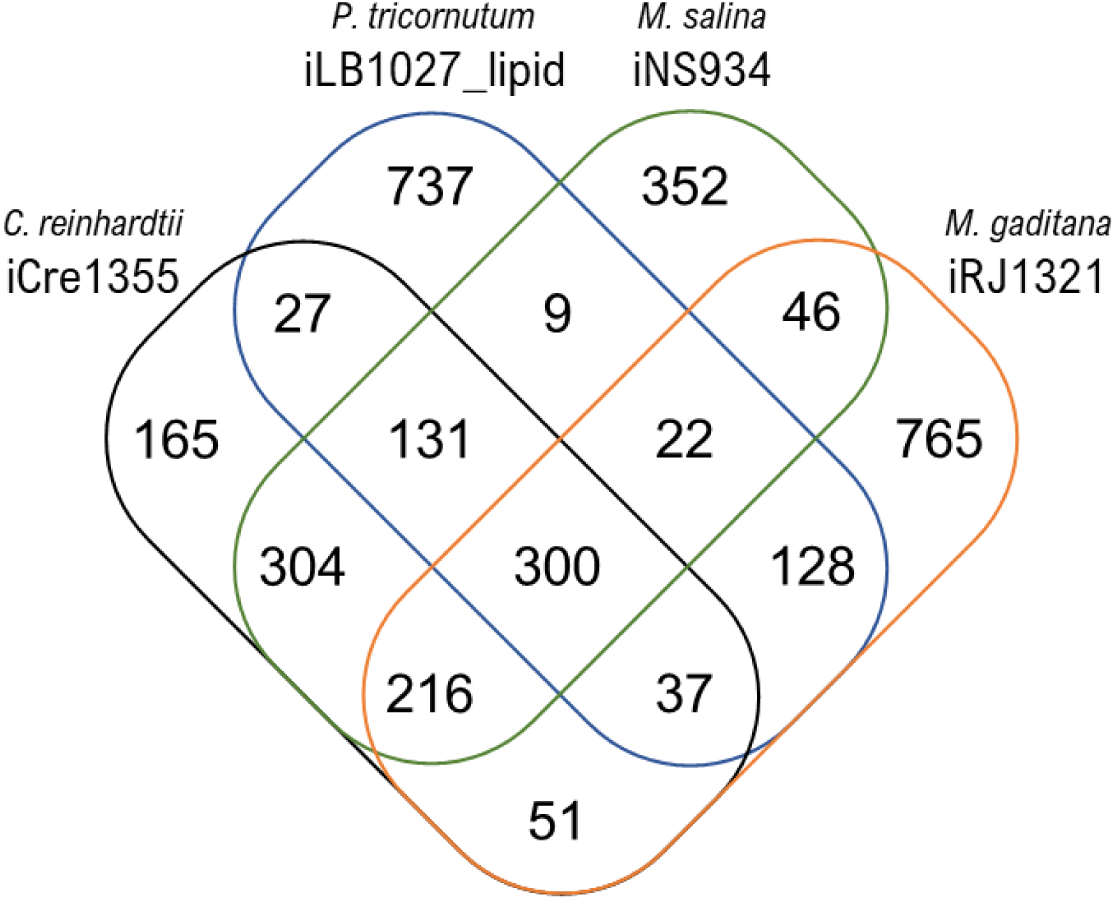
Relative scaffold GEM contributions to iSO1949 reactions, after automatic GEM construction and manual curation, based on reaction IDs.

### 3.3 Light-dependent biomass composition

The iSO1949_N.oceanica GEM differentiates growth of high-light and low-light acclimated *N. oceanica*, by adding light-dependent biomass synthesis reactions of high light conditions (HL, I_ph,0_ of 670 µmol.m^-2^.s^-1^, q_ph,abs_= 69.3 mmol_ph_.gDW^-1^.h^-1^) and low light conditions (LL, I_ph,0_ of 200 µmol.m^-2^.s^-1^, q_ph,abs_= 20.8 mmol_ph_.gDW^-1^.h^-1^). Biomass compositions were obtained from cultures of the lipidomics study on *N. oceanica* acclimated to varying light gradients and ingoing light intensities (Ferrer-Ledo et al., 2025).

The biomass reactions for HL and LL conditions consist of lipid, protein, carbohydrate, pigment and nucleotide fractions. Pigment and lipid species compositions were published previously (Ferrer-Ledo et al., 2025), and supplemented by carbohydrate and protein content analysis on the same triplicate biomass samples from all reactor conditions (Data S5). We observed no variation in carbohydrate content between HL and LL samples, which varied between 0.111 and 0.144 g/gDW. In other conditions, we observed only an increase in soluble carbohydrates to 0.163±0.021 and 0.171±0.016 g/gDW at I_ph,0_ of 200 µmol_ph_.m^-2^.s^-1^ and dilution rates of 0.025 h^-1^ and 0.031 h^-1^, which are more dilute conditoins than those used in the LL biomass dataset. Detected total protein contents varied between 0.261 and 0.354 g/gDW. A decrease in total protein composition could be observed in light inhibiting conditions only including the HL conditions, above net light absorption rates of q_ph,abs_ of 59 mmol_ph_.gDW^-1^.h^-1^. A clear decrease in protein content was observed at cyclostats with I_ph,0_ of 1550µmol_ph_.m^-2^.s^-1^ which is outside the range of the current study.

The total biomass content ranges from 0.65 to 0.68 g gDW^-1^. Part of the missing mass could be accounted for by ash content which could be up to 3-4% of the dry weight (*Nannochloropsis* spp., (Raso et al., 2012)), and by remaining moisture after freeze drying.

### 3.4 An extended pipeline for microalgal protein localization predictions

In this work, we extended on a previously developed pipeline (Levering et al., 2016; Sunaga et al., 2014) to predict the subcellular location of all annotated proteins. Our pipeline supplements the existing one with ASAFind for secondary plastids of red algal origin and with DeepLoc2.0 (Data S1). Although specific microalgal localization prediction tools exist for plants and green microalgae (PredAlgo (Tardif et al., 2012), TargetP (Armenteros et al., 2019)), protein localization in *Nannochloropsis* is complicated by its complex cellular structure. Heterokont algae evolved through double endosymbiosis, resulting in a 4-layer plastidic membrane merged with the ER. The specific origin of the double membranes varies per heterokont and has recently been proposed to deviate for *Nannochloropsis* compared to haptophytes and diatoms, which further complicates the prediction of subcellular localization of the proteins (Guo et al., 2019). The additional lack of validated protein localizations in double algal endosymbionts and consequent lack of training material for protein prediction tools reduces the predictive power of such tools for *Nannochloropsis*. In this work, we included specific heterokont tools HECTAR (Gschloessl et al., 2008) and ASAFind (Gruber et al., 2025, 2015). Although neither tool has been trained on *Nannochloropsis* sequences, ASAFind can sensitively predict the double signal-peptide with ‘ASAFAP’ motif needed for heterokont membranes and HECTAR has been shown to have a low false positive rate for *Nannochloropsis* mitochondrial and plastid sequences (Gruber et al., 2015; Vieler et al., 2012). Moreover, ASAFind 2 additionally predicts localizations into the periplastidial compartment (PPC). The PPC is located between the second and third membrane around the plastid, originally corresponding with the former cytosol of the red algal endosymbiont (Guo et al., 2019). The PPC forms a major site of transport between the plastid and other organelles, occasionally still housing several metabolic processes on its own (Ewe et al., 2018; Moog et al., 2011; Nonoyama et al., 2019). As is the case for most eustigmatophyceae, the PPC of *Nannochloropsis* has so far remained little explored. The combination of our predictions within the system-based perspective of the model might form the basis from which new insights can be obtained on the role of the PPC in *Nannochloropsis*.

We compared predictions of protein localization of the extended pipeline, to the initial pipeline developed by Levering et al. (Levering et al., 2016) Comparison between the pipelines shows that through our pipeline an increased number of nuclear genome encoded proteins locates to the nucleus, chloroplast and mitochondria, at the cost of cytosolic and peroxisomal localizations (Figure 4, Data S1). By using DeepLoc, proteins that previously remained unclassified and were therefore allocated to the cytosol are now assigned to mitochondria. The alternatively predicted plastidic proteins (1215 of 1942) would alternatively locate to the nucleus (215), mitochondria (4), ER (473), ER and/or cytoplasm (76), and cytoplasm (421) according to the pipeline of Levering et al. The shift in plastidic protein predictions can be attributed mainly to ASAFind and DeepLoc, in addition to reducing the weight of predictNL. Whereas the Levering pipeline considered all proteins with positive predictNLS hits to be located in the nucleus, we only assigned proteins to the nucleus when they scored higher on the nucleus than on other compartments. From the newly plastid categorized proteins, 334 were predicted with ASAFind, 625 with DeepLoc, 256 with both tools, and 67 were added through the adjustment of the nucleus scoring. Within the ASAFind2 predictions, 105 proteins were assigned to the PPC. Formerly, only 3 of these were predicted to be plastidic, 84 to be in the ER, and 16 to be in the nucleus by the former pipeline of Levering et al. However, it should be noted that the default PPC recognition pattern and threshold score are not optimized for *Nannochloropsis*. The large variation of localization predictions per prediction approach highlights the difficulty of predicting heterokont protein localization, and the requirement of consulting a range of localization tools.

**Figure 4.**
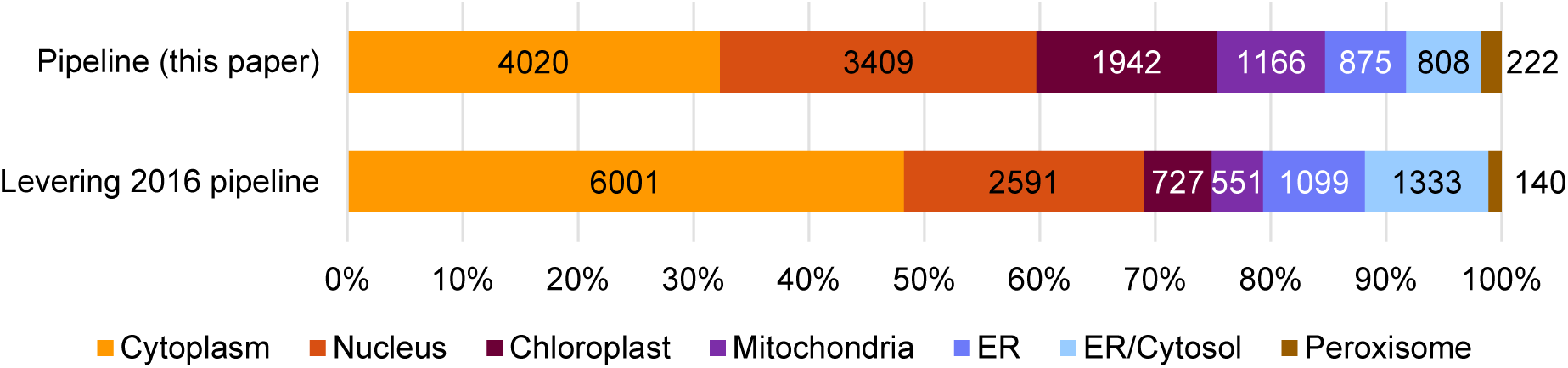
Predicted protein localization of the 12442 annotated proteins from the nuclear genome of *N. oceanica* IMET1, using both the prediction pipeline described in this paper and the pipeline described by Levering et al (Levering et al., 2016).

Often multiple genes are assigned to single reactions. We assessed whether the compartments reactions were assigned to match the predicted localization of their gene-associated protein products, also called Gene-Protein-Reaction (GPR). Most GEM reactions are linked to at least one enzyme that is predicted to be localized within the same compartment, even though some enzymes exhibit conflicting localizations, as highlighted in Appendix S1E. This discrepancy indicates both potential errors in enzyme assignments during model construction, as well as challenges in accurately determining enzyme localization. Moreover, the extended pipeline improves the match between model reaction locations and enzyme location predictions over the initial pipeline. Our findings indicate that it is important to consider discrepancies between predicted enzyme localization and assigned reaction localization when selecting genetic targets for metabolic engineering.

### 3.5 Curation of key metabolic pathways

During model construction and curation, special attention was paid to the curation of the core carbon metabolism. In this section we highlight key pathways unique to the iSO1949_N.oceanica GEM.

#### 3.5.1 Photosynthesis

Photosynthesis supplies and steers reducing power in the form of NADPH and ATP, driving CO_2_ fixation and subsequently the central carbon metabolism, including lipid synthesis. The linear electron transport consists of a series of reactions happening at the level of PSI, PSII, the cytochrome b6-f complex (CBFC), Ferredoxin/NADPH reductase (FNOR) and ATP synthase (ATPS). An overview of the photosynthesis reactions included in iSO1949_N.oceanica through orthology and gapfilling is indicated in Table 3. The model construction unveiled orthologs for all five linear electron transport reactions in the *M. gaditana* GEM and *N. oceanica*, whereas orthology for all reactions but PSII was found with the *M. salina* GEM. Orthology with the *C. reinhardtii* and *P. tricornutum* BEM was only found the FNOR reactions. The corresponding GPR to the orthologous photosynthesis reactions is presented in Data S6.

**Table 3.** Photosynthesis reactions in the iSO1949_N.oceanica GEM and scaffold GEMs. Grey cells indicate reactions included in iSO1949_N.oceanica through orthology (light grey) or through gapfilling (dark grey), white cells indicate reactions present in the scaffold GEMs that were associated with non-orthologous genes with *N. oceanica*. Empty white cells (-) indicate absence of the reaction in the respective scaffold GEM.

| Orthologue |  |  |  |  |
| --- | --- | --- | --- | --- |
| Gapfilled |  |  |  |  |
| iSO1949_N.oceanica | iLB1027_lipid | iRJ1321 | iNS934 | iCre1355 |
| *N. oceanica* | *P. tricornutum* | *M. gaditana* | *M. salina* | *C. reinhardtii* |
| Photosynthesis reactions |  |  |  |  |
| PSI_u | PSI_u | NanoG0619 | PSIred, PSIblue | PSIred, PSIblue |
| PSII_u | PSII_u | NanoG0616 | PSIIred, PSIIblue | PSIIred, PSIIblue |
| CBFC | CBFC_u | NanoG0617 | CBFC | CBFC |
| ATPSh | ATPS_h | NanoG0615 | ATPSh | ATPSh |
| FNOR_h, FNOR2_h | FNOR_h, FNOR2_h | NanoG0618 | FNORh | FNORh |
| Alternative electron flow |  |  |  |  |
| CEF_h | CEF_h | - | - | CEF |
| PTOX_h | PTOX_h | - | - | - |
| NDHPQR_h | NDHPQR_h | - | - | - |
| MEHLER_h | MEHLER_h | - | - | - |

Natural light access is dynamic, varying in intensity and exposure time on both short and long-time scales, ranging from seconds to days or months. Overexposure to light can saturate the linear electron flow (LEF) of photosynthesis, imposing a burden on the photosynthetic and cellular machinery. Alternative electron flow (AEF) enables algal cells to flexibly regulate light capture to maximize energy generation in any condition, without additional damage or costs to internal machinery (Bellan et al., 2020; Derks et al., 2015; Peltier et al., 2010). AEF can occur through cyclic electron flow (CEF), pseudo-cyclic electron flow (PCEF) and chloroplast-to-mitochondria electron flow (CMEF), collectively allowing for tunable processing of harvested electrons from the photosystems to generate ATP as discussed in detail by (Burlacot, 2023). Some of the scaffold GEMs incorporate AEF reactions, the most extensive being the *P. tricornutum* and *C. reinhardtii* GEMs. The cyclic electron flow reaction (CEF_h) and plastid terminal oxidase (PTOX) from *P. tricornutum*, orthologous to *N. oceanica*, were included in the model based on orthology (see Table 3). Additionally, the AEF Mehler reaction and NADH:plastoquinone reductase reactions from iLB1027, for which no orthologous genes were found, were also included in the model. The GEMs of *M. salina* and *C. reinhardtii* explicitly differentiate between photons with different wavelengths in the photosynthesis reactions. Since our dataset does not include light spectrum analysis, we considered non-wavelength-specific photon supply only and remapped wavelength-specific reactions to convert the generic ‘photon’ metabolite (see Data S6).

#### 3.5.2 Lipid biosynthesis

We systematically curated lipid biosynthesis under LL and HL, focusing on polyunsaturated fatty acid (PUFA) synthesis. In this section, we describe the key steps of fatty acid (FA) and PUFA biosynthesis, including the role of polar lipid classes in the desaturation process. Next, we compare how the orthology-based annotation of the lipid biosynthesis pathway compares to the genome annotation of the biosynthetic enzymes. The curated lipid metabolism is represented schematically in Figure 5. Overviews of the lipid species present in the *N. oceanica* and scaffold GEMs, as well as the detected lipid species *in vivo*, are provided in Data S7A-C.

**Figure 5.**
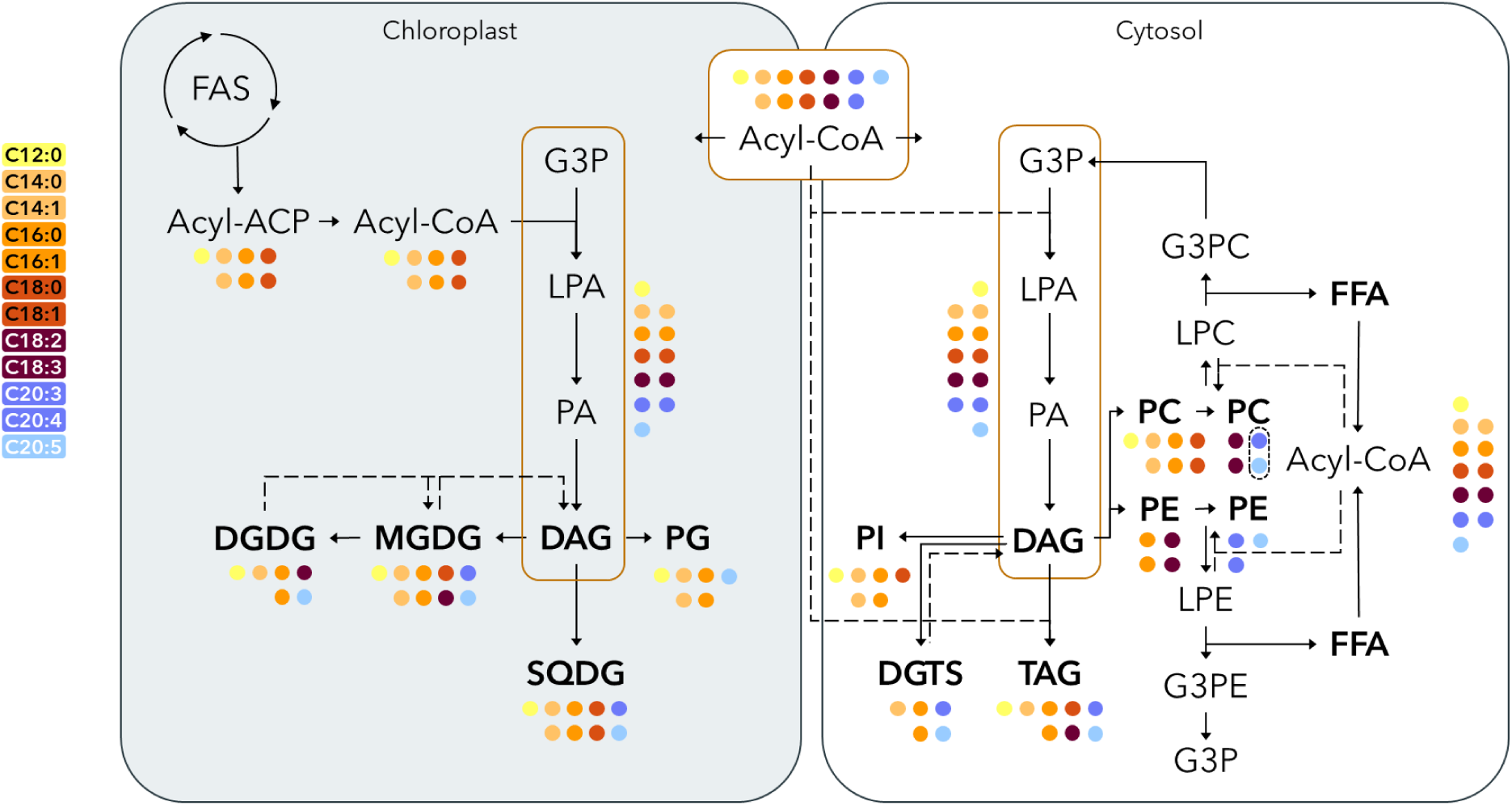
Schematic overview of the lipid metabolism described in the iSO1949_*N. oceanica* GEM based on lipidomic analysis, including degradation of lipid classes to allow for lipid class specific fatty acid desaturation. Colored dots represent the fatty acids present or used as intermediates in the respective lipid classes.

##### FA biosynthesis

iSO1949_N.oceanica represents algal lipid biosynthesis as initiated with fatty acid synthesis in the chloroplast, referred to as prokaryotic FA biosynthesis. Here, fatty acids are sequentially extended from acyl-CoA with malonyl-CoA (C2) into acyl-groups linked to acyl-carrier proteins (ACPs). The resulting plastidic acyl-ACPs have acyl-group lengths up to 12, 14, 16 or 18 carbon atoms. Acyl-ACPs can be directly used for plastidic lipid assembly or undergo further processing in the cytosol or ER via elongation and desaturation steps. The part of FA synthesis occurring outside of the chloroplast, known as eukaryotic FA biosynthesis, is responsible for producing long-chain mono- and polyunsaturated FAs (MUFAs and PUFAs) of up to 20 carbon atoms.

Conversion of free fatty acids into diacylglycerides (DAGs) and triacylglycerides (TAGs) occurs in the chloroplast and in the ER through the Kennedy pathway. Briefly, the FAs are activated by conversion into acyl-CoAs, followed by sequential double or triple attachment to a glycerol-3-phosphate backbone. Alternatively, diacyl intermediates from the Kennedy pathway, phosphatidic acid (PA) and DAG, can be re-processed further into a range of polar lipids by altering the polar headgroup. Moreover, polar lipids can transfer an acyl-group to DAG, resulting in acyl-independent TAG production.

##### PUFA biosynthesis

PUFA biosynthesis is a dynamic process of elongation and desaturation steps, taking place over multiple compartments and using several lipid classes as acyl-group desaturation sites. The exact compartmentalization of PUFA biosynthesis in *Nannochloropsis* species and the involved lipid classes are still under debate (Li et al., 2014; Xu, 2022).

Starting from C16:0 synthesized in the chloroplast, acyl-ACPs are elongated to C18:0 by palmitic acid elongase and desaturated to C18:1Δ9 by a plastidic stearoyl-ACP desaturase, followed by transfer out of the plastid to be processed further in the cytosol, ER and/or chloroplast ER (cER) (Dolch et al., 2017). Next, C18:1Δ9 is further desaturated by membrane-bound fatty acid dehydrases (FADs) on the 12^th^, and 6^th^ or 15^th^ carbon bonds by Δ12 and Δ6 or ω3 FAD respectively, resulting in C18:3ω6 or C18:3ω3 (Kaye et al., 2015; Poliner et al., 2018).

In *N. oceanica* C18 MUFA and PUFA species have been detected in high concentrations in phosphatidylcholine (PC) on the sn-2 position, suggesting that PC acts as scaffold for the desaturation of C18 (Schneider and Roessler, 1994; Vieler et al., 2012). Following desaturation C18:3ω6 and C18:3ω3 acyl-groups are released and elongated to C20:3ω6 and C20:3ω3 using a characterized Δ6 elongase, located in the cER (Shi et al., 2021). High contents of C20 PUFAs are detected in phosphatidylethanolamine (PE) and the betaine lipid diacylglyceryl-N,N,N-trimethylhomoserine (DGTS), suggesting that these lipid classes act as scaffolds for C20 desaturation. Desaturation of C20:3 to eicosapentaenoic acid C20:5ω3 (EPA) can start either using C20:3ω3 (Δ11,14,17) or C20:3ω6(Δ8,11,14) as substrate, catalyzed by Δ5 FAD and either Δ8 or ω3 FAD respectively.

Protein functionality of FADs and Δ6 elongase has been confirmed *in vivo* in *S. cerevisiae* and *N. oceanica* CCMP1779 (Poliner et al., 2018). In addition to desaturation on PC and PE/DGTS, studies have indicated that CoA-activated PUFAs are also substrates for desaturation (Δ5/6 acyl-CoA FADs), overcoming potential bottlenecks formed by reduced polar lipid availability (Khozin-Goldberg et al., 2011; Petrie et al., 2010).

##### Curation of eukaryotic PUFA biosynthesis in iSO1949_N.oceanica

We curated PUFA biosynthesis pathways as described above, based on the detected lipid species in *N. oceanica*. An overview of the detected lipid species and those used in the GEM biomass composition is provided in Data S7D-E. As can be seen in Data S7A, most lipid species of *N. oceanica* could be included in the draft GEM based on orthology with scaffold models. However, the scaffold GEMs varied widely in localization of lipid classes, resulting in an abundance of lipid species distributed over four compartments including the cytosol, chloroplast, ER and mitochondria. Moreover, whereas PC, PE and DGTS are reported as the main sites for PUFA desaturation in *N. oceanica*, different lipid classes sites are used by the scaffold models, occasionally lacking parts of the responsible pathways, leading to a partially inactive PUFA metabolism. In the following section, we assess FA desaturation as described by the scaffold models and indicate how PUFA synthesis was curated and adapted in the *N. oceanica* GEM to allow accurate simulation of PUFA synthesis and degradation.

Both biosynthesis and degradation of PC, PE and DGTS have to be present to allow for synthesis and transfer of PUFAs towards TAG. PC biosynthesis was present in all scaffold GEMs except for iCre1355. *C. reinhardtii* lacks PC, using DGTS instead as site for C18 elongation (Giroud et al., 1988). Although PC could be produced in the models iNS934 and iRJ1321, no desaturation or degradation steps were included in these GEMs. Only iLB1027 contained desaturation and degradation steps which took place on PC, allowing for release of the produced PUFAs. FAD reactions taking place on PC included desaturation of C18:1Δ9, C20:3ω3, and C22:5ω3 to C18:4ω3, C20:5ω3 and C22:6ω3 respectively. Only iLB1027 PC desaturase reactions were included in iSO1949_N.oceanica based on orthology, of which the C18:1Δ9 to C18:3ω6 desaturation reactions were kept after curation.

In contrast to PC, PE biosynthesis and degradation were present in all scaffolds although with inconsistencies. PE was used as FA elongation site only in the GEMs iNS934 of *M. salina* and iCre1355 of *C. reinhardtii*. Where iNS934 used plastidic PE to elongate C18:1Δ9 to C18:3 and C20:4 to C20:5 (isomers not specified), iCre1355 only elongated cytosolic PE C18:1Δ9 to C18:4ω3. *Chlamydomonas* is known to lack C20 PUFA’s (Li-Beisson et al., 2015).

The curation process led us to detect inconsistencies on the GEMs used as a scaffold, especially iNS934. iNS934 contains pathways for both cytosolic and plastidic PE biosynthesis, but degradation and elongation are only present in the plastid. Although the plastidic PE can be used as a site for plastidic PUFA synthesis, it cannot be used as a PE source for growth since the iNS934 biomass equations only contain cytosolic PE and any PE transfer between compartments is absent. Therefore, our analysis also produces suggestions to further improve those models. The PE FAD reactions from both iNS934 and iCre1355 were orthologous to *N. oceanica* and included in the iSO1949_N.oceanica draft. We moved the PE C20:4ω3 Δ5 FAD to the cytosol, and gapfilled C20:3 to C20:5 desaturation using the orthologous PC-specific Δ5 FAD and ω3 FAD reaction from *P. tricornutum* as template. Any other PE FAD reactions in the plastid or targeting C18 species were removed.

The lipid class DGTS was present in the GEMs of *M. salina*, *M. gaditana*, and *C. reinhardtii*. Both the *M. salina* and *C. reinhardtii* GEMs showed orthology for DGTS biosynthesis. However, the *M. salina* GEM included DGTS synthesis in the plastid similar to PE, even though the orthologous *N. oceanica* GPR is predicted to be ER and mitochondria localized. Only the *C. reinhardtii* GEM included reactions to further desaturate cytosolic C18:1 to C18:3 and C18:4ω3 PUFAs, although the synthesized PUFAs are not released indicated by the lack of DGTS degradation. *Chlamydomonas* does not require C18 PUFA transfer for further elongation or incorporation into other lipid classes (Li-Beisson et al., 2015), as is the mechanism in *Nannochloropsis*. The *C. reinhardtii* DGTS desaturation reactions were transferred to the iSO1949_N.oceanica draft through orthology. Reactions of FAD Δ5 and ω3 were edited to target C20:3ω6 and C20:4ω3/6 instead of C18 species. We noted that the GPR of ω3 FAD assigned through orthology (NO14G02790.1) matches the functional annotation from the NanDeSyn DB. Interestingly, the *P. tricornutum* GEM lacked the DGTS species entirely despite DGTS being a major lipid class in *Phaeodactylum* (Popko et al., 2016) and iLB1027 being especially curated on the *Phaeodactylum* lipid metabolism.

The scaffold GEMs contain partial pathways for synthesizing PUFAs but most do not represent the PUFA biosynthesis pathways of *N. oceanica*. The lipid classes PC, PE and DGTS are merely included in these GEMs as biomass components, not as intermediates in lipid synthesis. Instead, CoA-activated PUFA desaturation is the main mode of desaturation. Acyl-CoA targeting FAD and elongase reactions were present in all scaffolds besides iLB1027, covering the complete PUFA synthesis from C18:1Δ9 to C18:3ω3 for iCre1355 and to C20:5ω3 for the *Microchloropsis* GEMs. In addition, initially the complete acyl-CoA desaturation pathways converting C18:1Δ9 to C20:5ω3 were included in the *N. oceanica* draft GEM based on orthology, resulting in parallel PUFA synthesis pathways either using lipids as scaffolds or free acyl-CoA’s. Interestingly, FAD genes were included through orthology-based GEM construction which differed from functionally annotated FADs from the genome, suggesting possible isomers. An overview of all annotated FADs in the draft iSO1949_N.oceanica GEM is provided in Data S7E. Locus tags and the predicted localization of the associated reactions in the GEM are provided in Data S7E as well. In the final *N. oceanica* model we adapted PUFA desaturation as described in the paragraphs above.

##### Curation of plastidic PUFA biosynthesis in iSO1949_N.oceanica

Besides FA desaturation in the eukaryotic PUFA biosynthesis, several pathways for plastidic PUFA biosynthesis were partially provided to the iSO1949_N.oceanica draft as well that are not all native to *N. oceanica*. Main lipid classes in the plastid are galactolipids MGDG and DGDG, the phospholipid phosphatidylglycerol (PG) and sulfolipid sulfoquinovosyldiacylglycerol (SQDG), which are all synthesized in the chloroplast using DAG as substrate. The draft included desaturation reactions that used MGDG and DGDG as site for C16-specific Δ4, Δ7 and ω3 desaturation to C16:4ω3(4,7,10,13Z) owing to orthology with the *C. reinhardtii* GEM, in addition to C18:2(9,12Z)-specific Δ5 and ω3 desaturation to C18:3ω6(5,9,12Z) and C18:3ω3(9,12,15Z). Moreover, C18:2(9,12Z)-specific ω3 desaturation on SQDG was included from the *C. reinhardtii* GEM. Additionally, although plastidic PUFA biosynthesis is proposed to be the main mode of C16:3ω3 but also C16:3ω4 synthesis in *P. tricornutum* (Mühlroth et al., 2013), the C16:3ω4 species was detected in *N. oceanica* in the PC and TAG lipid classes only, suggesting that 16:3ω4 is formed outside the chloroplast from extra-plastidial 16:2ω7 by a Δ12 desaturase on the PC, similar to C18:3ω6 synthesis from 18:2ω9. Moreover, no plastidic FAD 12 has been annotated in *N. oceanica* that could provide precursors for plastidic C18:3ω6(5,9,12Z) formation. We transferred this PUFA biosynthesis on PC in the iSO1949_N.oceanica GEM.

Moreover, plastidic PG specific C16Δ3^t^ FAD was present in the draft iSO1949_N.oceanica GEM based on orthology with both *C. reinhardtii* and *P. tricornutum*. The PG C16Δ3^t^ FAD converts PG sn-2 C16:0 to PG sn-2 C16:1Δ3E and has been annotated in *N. oceanica* (Vieler et al., 2012). The C16:1Δ3^t^ in PG is involved in stabilizing PSII (Kruse et al., 2000). Besides in PG, C16:1Δ3E has been detected in SQDG as well by Schneider and Roessler (Schneider and Roessler, 1994) suggesting miscellaneous specificity of the C16:1Δ3E FAD and possible function of SQDG as an alternative to PG in *N. oceanica* (Bolik et al., 2022). Although only the PG C16:1Δ9 lipid was included in the biomass equations of iSO1949_N.oceanica, since our detection methods cannot distinguish between C16:1(9Z) and (3Z) isomers, this lipid metabolite can be considered representative for both the Δ3E and Δ9 isoforms.

##### PUFA shuttling between compartments

Although in *N. oceanica* PUFA biosynthesis occurs via eukaryotic FA synthesis, the highest ratio of EPA is found in polar galactolipids of the plastid, suggesting EPA shuttling between compartments (Schneider and Roessler, 1994), (Sukenik et al., 1989). EPA transfer from the ER to the plastid has been investigated in *Porphyridium* (Khozin et al., 1997) and *Chromysta* (Mühlroth et al., 2013), but the involved EPA lipid class carriers remain unclear. Proposed intermediates are DGTS (Vieler et al., 2012), PE (Meng et al., 2017), and DAG (Han et al., 2017), the latter possibly derived from catabolized phospholipids, although lysis of acyl-groups to acyl-CoA followed by transport in the form of free acyl-CoA has been considered as well. However, DGTS conversion to putative lipid transfer intermediates DAG or TAG is unlikely due to the absence of enzymes capable of breaking the ester bond between the glycerol backbone and the headgroup, and the lack of DAG species with a matching acyl-group composition (Hoffmann and Shachar-Hill, 2023). Instead, DGTS has been proposed as an alternative EPA sink, similar to and competing with MGDG in this role (Murakami et al., 2018). ASAFind 2.0 predictions for PPC localized enzymes included three enzymes involved in the lipid metabolism, including a glycerol-3-phosphate dehydrogenase (GPDH, NO18G01950.1), fatty acid hydroxylase (NO03G03660.1), and NO20G02200.1 which is associated in the GEM with a PE-specific phospholipase reactions. These predictions are the first indications for *Nannochloropsis* that the PPC might play a role in lipid metabolism, possibly during the severe lipid trafficking and re-shuffling and high accumulation of TAG during nitrogen starvation (Janssen et al., 2020, 2019).

##### Additional curation of minor fatty acid species

C12 and C14:1Δ9 species were detected in several lipid classes, but are often not reported in literature (Ferrer-Ledo et al., 2025). Other *N. oceanica* studies (Liu et al., 2013; Meng et al., 2017) confirm the presence of these FA species, albeit at low percentages. The detected C12 and C14:1Δ9 containing lipids were added manually to the GEM and can be included or excluded in the biomass reactions according to the users’ preference.

Glycerolipids are known to be produced by plants across compartments, in either the plastid or endoplasmic reticulum (Benning, 2009). In the iSO1949_N.oceanica GEM we did not separate and restrict the localization of these pathways, because the localization predictions of lipid biosynthesis enzymes are still unclear. Therefore, to prevent lipid reactions from not having access to relevant substrates and energy carriers, we curated the eukaryotic part of lipid synthesis to be located in the cytosol and did not include the endoplasmic reticulum compartment.

#### 3.5.3 Carotenoid biosynthesis

Pigments play an essential physiological role as antennae for harvesting protons in photosynthesis and in energy dissipation through the xanthophyll cycle. Although microalgal pigments are industrially relevant products (Lubián et al., 2000), pigments make up a relatively small fraction of the biomass in microalgae including *Nannochloropsis*, and are therefore often overlooked in metabolic models and databases. In addition, pigment composition largely varies between algal species. Here, we curated the carotenoid pathway in detail to the physiology of *N. oceanica* and specifically considered the usage of reducing equivalents.

*Nannochloropsis* makes use of chlorophyll a in addition to the carotenoids antheraxanthin, zeaxanthin and violaxanthin in the light harvesting complexes. Accessory pigments are astaxanthin, β-carotene, canthaxanthin and vaucheriaxanthin, although they are present at low concentrations. An overview of the pigments detected and curated for iSO1949_N.oceanica and those present in the scaffold GEMs is provided in Table 4. Our measurements of *N. oceanica* pigments detected carotenoids and chlorophylls, including the less reported and less abundant vaucheriaxanthin and neoxanthin. Vaucheriaxanthin contents were comparable to previous reports (Amaral et al., 2021) and (Borowitzka et al., 2016), its exact biosynthesis mechanism has not yet been reported (Basso et al., 2014; Keşan et al., 2016; Lubián et al., 2000). Neoxanthin is an intermediate for vaucheriaxanthin synthesis, and has been detected in trace amounts in *N. oceanica* (Liu et al., 2022) but was not present above detection thresholds in our measurements. We included biosynthesis reactions to also produce these less abundant metabolites.

**Table 4.** Pigments present in the iSO1949_N.oceanica GEM and scaffold GEMs. Pigments included in the iSO1949_N.oceanica GEM are indicated by the respective metabolite identifiers. For the scaffold GEMs, BM indicates pigments included into the biomass reaction, and ‘Metabolite’ indicates pigment only present as intermediate metabolite. Pigments indicated with asterisks (*) are present in the scaffold models, but absent from *N. oceanica*.

| Pigment | iSO1949_N.oceanica<br>Metabolite ID | iNS934 | iRJ1321 | iLB1034 | iCre1355 |
| --- | --- | --- | --- | --- | --- |
| Chlorophyll a | chla_h | BM | BM | BM | BM |
| Chlorophyll b* |  | Metabolite |  |  | BM |
| a-Carotene | acaro_h | BM | Metabolite | Metabolite | BM |
| b-Carotene | caro_u / h | Metabolite | BM | Metabolite | BM |
| Canthaxanthin | canxan_u |  |  |  |  |
| Astaxanthin | astxan_u |  |  |  |  |
| Zeaxanthin | zeax_u / h | BM | BM | Metabolite |  |
| Antheraxanthin | anxan_u / h | BM | Metabolite | Metabolite | BM |
| Violaxanthin | vioxan_u / h | BM | BM | Metabolite | BM |
| Vaucherixanthin | vauxan_u |  |  |  |  |
| Neoxanthin | neoxan_u | BM |  | Metabolite | BM |
| Lutein | lut_u | BM |  |  | BM |
| Loroxanthin* |  | BM |  |  | BM |
| Fucoxanthin* |  |  |  | BM |  |
| Diodinoxanthin* |  |  |  | BM |  |
| Chlorophyll C1/2 * |  |  |  | BM |  |

The initial draft GEM included the complete initial pigment biosynthesis pathway up to phytoene. Parallel pathways were present leading from phytoene to β-carotene through either neurosporene or tetra-cis-lycopene. These pathways markedly differ in the usage of reducing equivalents usage. Formation of carotenoids downstream from β-carotene was initially localized to both the thylakoid and chloroplast (stroma) compartments. The xanthophyll cycle involving zeaxanthin, antheraxanthin and violaxanthin was associated to orthologous genes in all 4 scaffold GEMs, emphasizing preservation of this mechanism between microalgae. The iRJ1321 and iLB1027 scaffold GEMs localized this pathway in the chloroplast, iNS934 and iCre1355 scaffolds used the thylakoid compartment instead, resulting in mixed localization in the draft GEM.

We unified the inherited carotenoid pathways into the thylakoid and GPR associations were combined when available. Cofactors were considered to be taken from the chloroplast reducing equivalent pool to ensure energy coupling between pigment synthesis and the central metabolism. Additional reactions were added for neoxanthin, vaucheriaxanthin, astaxanthin and canthaxanthin biosynthesis. Although no orthologues were found for neoxanthin formation, a violaxanthin de-epoxidase-like protein (VDL) has been investigated in *N. oceanica* CCMP1779 (Dautermann et al., 2020). Based on homology, the reaction was added to the draft associated with gene NO44G00920.1 (Data S8B). Other carotenoids added to the model are astaxanthin and canthaxanthin, which were only detected under light-inhibiting conditions in the reactor studies (Ferrer-Ledo et al., 2025). Biosynthesis reactions for astaxanthin and canthaxanthin were added from *C. reinhardtii*, based on (Perozeni et al., 2020; Zhong et al., 2011) and the MetaCyc database. Model curation of the carotenoid biosynthesis pathway was complemented with experimental data from Liu et al. (Liu et al., 2022) (Data S8A).

The initial model contained three alternative lycopene biosynthesis pathways from phytoene, see Figure 6. Two of these pathways are known to be present in bacteria and fungi (Estrada et al., 2008; Hausmann and Sandmann, 2000; Schaub et al., 2012) while the other has been found in plant and cyanobacteria. Both multistep pathways use the enzymes phytoene desaturase (PDS) and ζ-carotene desaturase (ZDS) in addition to neurosporene oxidoreductase (NOR) in the bacterial/fungal pathway and ζ-carotene *cis-trans* isomerase (Z-ISO) and carotene *cis-trans* isomerase (CRTISO) in the cyanobacteria/plant pathway. The draft GEM inherited the single-enzyme reaction from the *P. tricornutum* GEM, and both multistep-pathways from the *M. gaditana* and *C. reinhardtii* GEM. The *M. salina* GEM contained the plant pathway only. The PDS and ZDS reactions both require the electron carrier plastoquinone. Reduction by PDS of 2 plastoquinones has been confirmed for in *C. reinhardtii*, with similar functioning shown for ZDS in *Arabidopsis* (Norris et al., 1995; Vila et al., 2008). Although plastoquinones are consumed in all three pathways, only two pathways result in plastoquinol production. The other pathway has no net plastoquinol production but use O_2_ as cosubstrate instead in the ZDS and NOR reactions (Figure 6) in a similar manner to the PTOX reaction.

**Figure 6.**
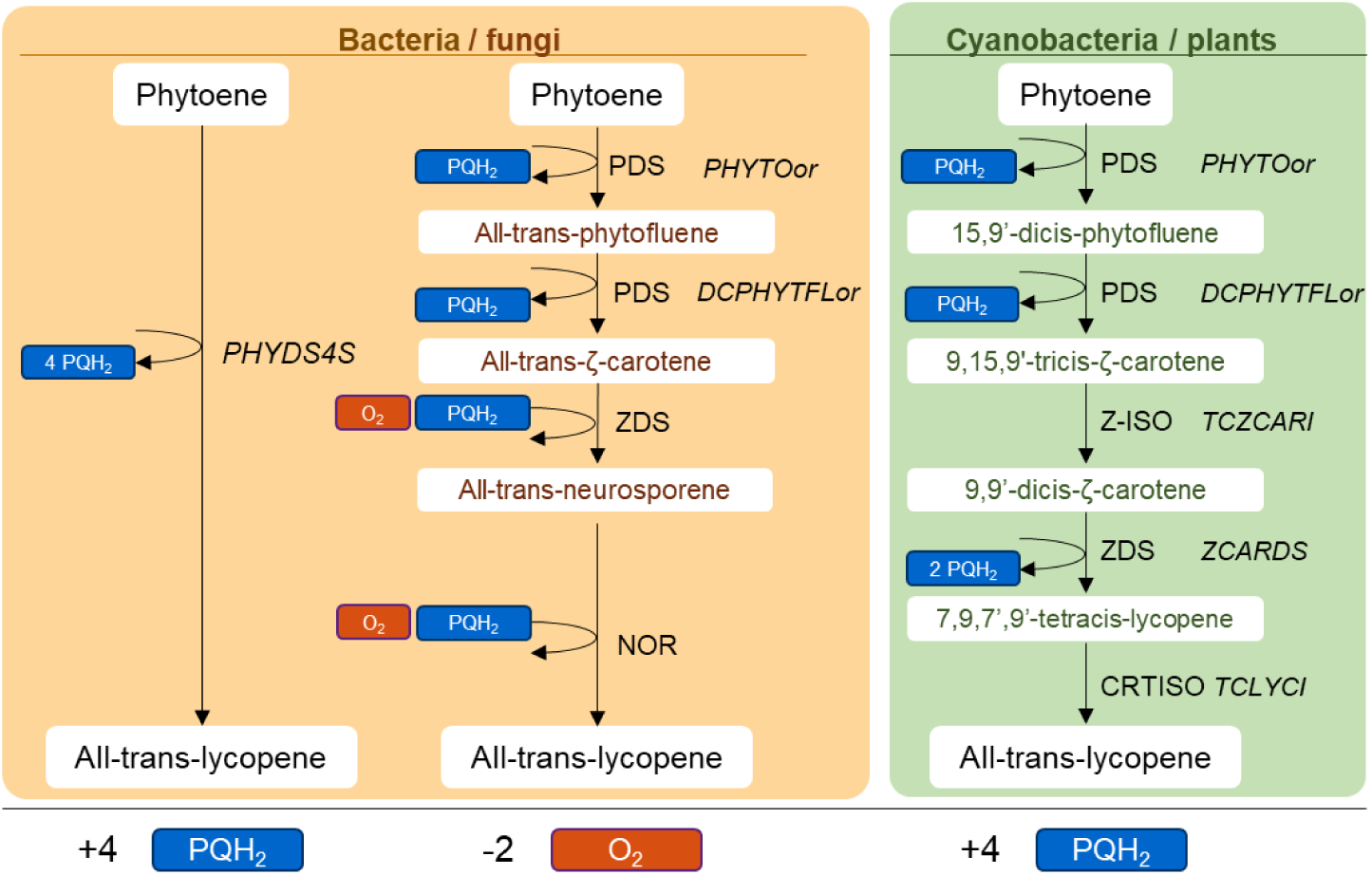
Demonstration of parallel pathways with differing cofactor usage, resulting from GEM construction. Here we show alternative pathways of lycopene biosynthesis from phytoene with varying cofactor usage, as present in the *N. oceanica* model. Text in boxes represents metabolites, text outside of the boxes in standard and cursive font indicates the corresponding enzyme and GEM reaction ID respectively. PDS - phytoene desaturase, ZDS - ζ-carotene desaturase, NOR - neurosporene oxidoreductase, Z-ISO - ζ-carotene cis-trans isomerase, and CRTISO - carotene cis-trans isomerase.

Inheritance of the three lycopene pathways highlights the impact that annotation and model curation have on metabolic modeling. Due to the different net consumption of reducing power, exclusively the least costly pathway in terms of plastoquinol will be used by the GEM in simulations, potentially impacting GEM predictions. Consequently, the predictions generated by the GEM may not identify genes associated with the pathways that require higher cofactor usage, despite their potential relevance to metabolic engineering.

### 3.6 Modelling maintenance requirements using Photosynthesis-Irradiance curves

More than accurate representation of the metabolic pathways, GEM performance is also determined by accurate constraints of energy requirement for cellular maintenance. Cellular maintenance in the GEM is represented by growth associated maintenance (GAM) and non-growth associated maintenance (NGAM). GAM describes the additional energy needed to replicate biomass and is included in the biomass equation as additional ATP requirement. NGAM on the other hand, is the energy required to sustain 1 gram of biomass per hour and is included in the GEM as an ATP consuming reaction that is forced to carry a positive flux. The iSO1949_N.oceanica maintenance rates were constrained using experimentally determined respiration measurements of high light (HL) and low light (LL) acclimated cultures.

NGAM rates for *N. oceanica* were determined using photosynthesis-irradiance (PI) curves for LL and HL acclimated phenotypes, shown in Figure 7. PI-curves represent the short-term photosynthetic response to varying surface irradiance levels, or light intensities, in the form of O_2_ and CO_2_ conversion rates. We normalized this surface irradiance by the absorbed light throughout the culture by biomass, making it compatible with model simulations. PI-curves can provide a ‘metabolic snapshot’, giving some insight into the light-driven respiratory capabilities, or energetic needs, of a given phenotype without long-term acclimation of the metabolism.

**Figure 7.**
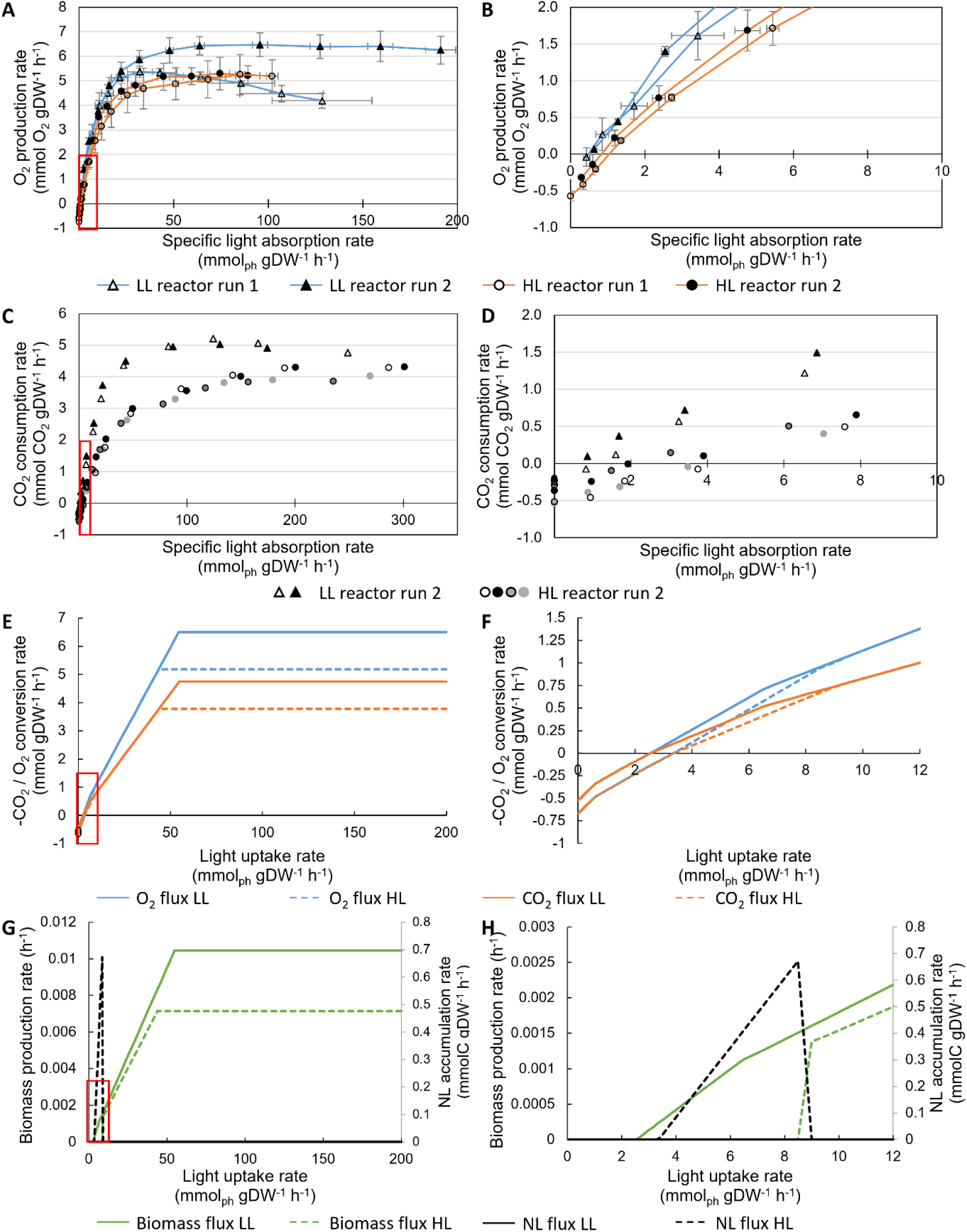
PI-curves N. oceanica acclimated to low and high light. Panels B, D, F, and H are zoomed in versions of the red outlined parts of panels A, C, E, and G respectively. Panels A-B: measured oxygen production rates of individual samples of two bioreactors per light setting (LL, blue, n=3-4, HL, orange, n=4). Panels C-D: measured CO2 consumption rates of individual samples of one LL and one HL reactor (LL, triangles, n=2, HL, circles, n=4). Panels E-F: simulated CO2 consumption (orange) and O_2_ production (blue) rates at increasing light intensities, simulated using LL and HL specific biomass compositions and NGAM rates and by optimizing for maximal CO2 uptake. LL is indicated by solid lines, HL by dashed lines. Panels G-H, biomass production rate and C-mol normalized neutral lipid accumulation rate. LL is indicated by solid lines, HL by dashed lines. One of the LL bioreactors showed early light inhibition and was excluded for calculating maximum O_2_ production (panel A, white triangles).

Oxygen uptake rates measured in dark conditions can be considered representative for the minimum energy required to sustain cells, as discussed by Kliphuis et al. (Kliphuis et al., 2011), and have been used in a similar manner to constrain NGAM in a *Synechococcus* GEM (Broddrick et al., 2019). We determined O_2_ uptake rates of 0.310±0.078 and 0.419±0.114 mmolO_2_ gDW^-1^ h^-1^ respectively, for *N. oceanica* acclimated to LL and HL, detected after 30 minutes of dark acclimation ^(^Table 5^)^. These rates are comparable to previously determined rates of *Chlorella kessleri, C. reinhardtii*, and *N. oleabundans*, which were 0.36, 0.53, and 0.47 mmol O_2_ gDW^-1^ h^-1^ respectively (Kliphuis et al., 2012, 2011; Klok et al., 2013). Alternatively, NGAM rates can be obtained through chemostat experiments as described by (Thiele and Palsson, 2010). Graphs relating to growth rates and substrate requirement can be extrapolated to find the minimum substrate requirement associated to the maintenance rate. Recent chemostat studies in *Nannochloropsis sp.* and *M. salina* determined the minimum photon uptake rate requirements. Assuming an 8:1 photon:O_2_ ratio, minimal respiration requirements of 0.24 and 0.95 mmolO_2_ gDW^-1^ h^-1^ are found for *Nannochloropsis* sp. and *M. salina* respectively (Barten et al., 2022; Sforza et al., 2015).

**Table 5.** Parameters determined using O2 and CO2 photosynthesis-irradiance curves (PI-curves) for LL and HL *acclimated N. oceanica* cultures. SD represents standard deviation of n=3 and n = 4 for the LL replicate reactors, and n = 4 for the HL replicate reactors.

|  |  | Respiration rate / NGAM |  | Maximum O<sub>2</sub>/CO<sub>2</sub> conversion |  | Light compensation point |  |
| --- | --- | --- | --- | --- | --- | --- | --- |
|  |  | mmol gDW<sup>-1</sup> h<sup>-1</sup> |  | mmol gDW<sup>-1</sup> h<sup>-1</sup> |  | mmol<sub>ph</sub> gDW<sup>-1</sup> h<sup>-1</sup> |  |
|  |  | Average | SD | Average | SD | Average | SD |
| O<sub>2</sub> | LL Replicate reactor 1 | -0.296 | 0.078 | 5.39 | 0.20 | 0.54 | 0.17 |
|  | LL Replicate reactor 2 | -0.316 | 0.086 | 6.50 | 0.39 | 0.52 | 0.06 |
|  | Average LL | -0.310 | 0.078 | 6.02 | 0.66 | 0.53 | 0.07 |
|  | HL Replicate reactor 1 | -0.344 | 0.100 | 5.21 | 0.40 | 0.73 | 0.22 |
|  | HL Replicate reactor 2 | -0.493 | 0.073 | 5.15 | 0.59 | 1.17 | 0.07 |
|  | Average HL | 0.419 | 0.114 | 5.18 | 0.48 | 0.98 | 0.27 |
| CO<sub>2</sub> | LL | 0.225 | 0.049 | -5.12 | 0.12 | 0.86 | 0.41 |
|  | HL | 0.422 | 0.127 | -4.12 | 0.22 | 3.22 | 0.92 |

Assuming all O_2_ taken up is used for ATP generation through linear electron transport in a 1:3 molar ratio, the *N. oceanica* O_2_ uptake rates can be used to determine NGAM rates for the *N. oceanica* GEM. We found NGAM rates of 0.930 and 1.256 mmol ATP gDW^-1^ h^-1^ for the LL and HL biomass equations respectively, which are comparable to those of the scaffold GEMs, which varied between 1.00 and 2.85 mmol ATP gDW^-1^ h^-1^ for *P. tricornutum*, *M. gaditana*, and *C. reinhardtii*. The *M. salina* GEM did not report NGAM constraints.

### 3.7 Influence of light-acclimation constraints on photosynthesis and growth

We assessed the ability of the GEM to simulate photosynthesis through comparison of model predictions and measured PI-curves. Moreover, we assessed whether the GEM could replicate the effect of light-acclimation on photosynthesis by simulating PI curves using the LL and HL acclimation-dependent NGAM rates, maximal O_2_ production rates and biomass equations.

PI-curve measurements for LL and HL acclimated phenotypes and the GEM PI-curve simulations are shown in Figure 7 and detailed in Data S5. In order to simulate PI-curves, we included the following constraints. First, NGAM and O_2_ exchange constraints were set based on the previously discussed dark respiration rates and maximum O_2_ production rates shown in Figure 7A-B (Table 5). Second, to represent usage of storage compounds in the dark, cytosolic glucose uptake was allowed when simulating light conditions below the light compensation point. As a final constraint the model was allowed to store the fixated carbon either in biomass or in neutral lipid pool, and either high light or low light acclimation-specific compositions.

The GEM was able to simulate the maximum CO_2_ uptake rate comparable to the experimental values (Figure 7C-D and Figure 7E-F), when constrained with the measured maximum O_2_ production rate. The ability to replicate maximal CO_2_ fixation indicates that the GEM is energy-balanced, as it connects the energy generation of photosynthesis to carbon fixation in the correct ratio without interference of artificial ATP generating loops. Given the limited constraints for phototrophic algae on substrate and product exchange rates, correct connection between O_2_ and CO_2_ exchange rates is essential to ensure accurate predictions of metabolic behavior.

Both the experimental and simulated PI-curve show increased O_2_ production and CO_2_ consumption rates for the LL phenotype compared to the HL phenotype (Figure 7A-F). In LL conditions, microalgae adapted their cellular machinery to increase light capture, thus resulting in an increased photosynthesis rate and optimal carbon fixation yields, compared to HL conditions. On the other hand, under prolonged HL conditions, photodamage causes a reduction of the photosynthetic machinery and membranes and machinery, resulting in decreased O_2_ production compared to LL conditions. Moreover, under HL captured energy is re-directed to the synthesis of high-energy containing metabolites such as lipids, which results in less energy and carbon allocated for growth. The decreased growth rate of 38% compared to a decrease of maximal CO_2_ fixation of 20% shows that the GEM allocates the additional energy cost of the HL biomass as higher lipid energy-dense contents (Figure 7G). On the other hand, the model simulations indicate similar rate of change in CO_2_ consumption and O_2_ production for both the LL and HL modes, with respect to changes in light uptake (slope of the curves in Figure ^7^E). This contradicts the experimental data where we observe differences in the slope of the corresponding curves between the two acclimation states (Figure 7A, C). This is likely due to changes in light absorption kinetics by pigments that are difficult to capture by the constraint-based approach here deployed. Moreover, the model assumes complete use of the provided photons for activating photosynthesis, whereas only a selection of protons has the compatible wavelength.

The GEM was able to simulate comparable light compensation points (I_ph,c_) of LL and HL conditions, and simulated increased I_ph,c_ for HL compared to LL, see (Figure 7F). In the GEM, the I_ph,c_ is determined by constraints on the NGAM and by the cyclic electron flow (CEF) (Data S5C). A higher NGAM, as is the case for the HL phenotype, requires an increased amount of ATP from photosynthesis, thereby increasing the photon supply threshold for reaching net O_2_ production, whereas the CEF requires photons to be used by PSI thereby competing for photons with the oxygen producing PSII.

While higher experimental I_ph,c_ values were found for the CO_2_ curves compared to the O_2_ curves, the GEM simulated these to be the same. The higher *in vivo* I_ph,c_ for the O_2_ production compared to CO_2_ consumption indicates that the ratio between O_2_ and CO_2_ are different between respiration and carbon fixation through photosynthesis.

## 4 General discussion

The iSO1949_N.oceanica GEM provides a knowledge base of the metabolism of *N. oceanica.* The GEM captures an extensive collection of genes coupled to their metabolic functions and to the subcellular location of their protein products. The consistency of the GEM here presented has been extensively tested. The curation process has ensured the model fulfils consistency requirements, for instance no biomass or other products can be obtained without energy and a carbon source as the model cannot generate ATP or reducing equivalents without light or the use of a storage C-source. This is a minimal consistency requirement that models of phototrophic organisms should fulfil, as included in GEM consistency requirements by Pham et al. (Pham, 2021). The simulations shown in Figure 7 indicate that the model correctly captures the light response phenotype of the organism. Nevertheless, additional validations and adaptations are needed to further refine the GEM and reduce redundancy.

Experimental data of enzymatic functions, localizations, expression levels and constraints under varying environments can contribute to further tailoring of the GEM. *N. oceanica* is known to contain a larger abundance of biosynthetic lipid gene isomers compared to other model green algae, for example (Wang et al., 2014), which vary in their role and localization in the cell (Nobusawa et al., 2017; Zienkiewicz et al., 2017). However, genome-based metabolic models are not able to discriminate in enzyme isomer roles based on gene annotation, therefore requiring additional curation, and eventually, experimental biochemical validation. Moreover, the orthology-based reconstruction provided additional putative isomers, and localization tags provided alternative putative localizations, all of which provide leads for subsequent experimental studies.

In addition, pathways such as lipid degradation pathways, can be refined in the model. These have not been the point of focus of this study and therefore have not been curated. Furthermore, alternative reaction localizations can be assessed by relocating metabolic reactions within the GEM based on the enzyme localization information. Simulation of such alternative pathways can provide new insights and pinpoint knowledge gaps on microalgal metabolism, and it can also increase our ability to predict localizations.

The model can be further constrained using existing omics data (Yurkovich and Palsson, 2018), limiting the active reactions to the subset associated to the expressed genes, thereby reducing GEM redundancy for simulations. Comparison with low-light and high-light acclimated *N. oceanica* transcriptomics specifically in combination with the acclimated photophysiology, will allow simulation of pathways underlying the light-dependent phenotypes. In turn, less redundant simulation of the light-dependent phenotype also allows for clearer assessment of genetic engineering effects under such light conditions.

Our PI-curve simulations showed that the GEM can simulate short-term light response from long-term light acclimated phenotypes. These data are obtained using the BOM, an accessible tool widely used in the microalgae field. However, although the GEM can provide insights into the effect of constant variables such as light absorption, nutrient availability and photophysiology, kinetic effects cannot be predicted yet, such as photoinhibition, photo acclimation through non-photochemical quenching, and photorespiration. These kinetic processes take place on a timescale of minutes to hours and affect light-harvesting in dynamic reactor conditions. Kinetic models have been developed that can translate such effects to modulation of photosystems and of the alternative electron flow (Saadat et al., 2021). Such models can be combined with the GEM, which can provide snapshots of the underlying metabolism given provided photosystem constraints.

Although steps can be taken to further extend the model, iSO1949_N.oceanica is ready to be applied for performing target predictions for metabolic engineering. The GEM can be applied to predict carbon partitioning related to *N. oceanica* specific biomass production, pigment and lipid biosynthesis pathways. Additionally, target enzymes can be predicted for increasing neutral lipid productivity or tailoring lipid class compositions for example. Moreover, the light-acclimated constraints provide users with an approach to assess the implications of metabolic engineering on the metabolism under the variable light conditions present in photobioreactors.

Although GEMs have been used increasingly to describe microalgal metabolism of model organisms, few exist for industrially relevant microalgae. In this study, we developed the first genome-scale metabolic model of the oleaginous microalga *N. oceanica* in a traceable way, tailored in detail with biochemical and photophysiology constraints obtained under steady-state conditions to high light and low light acclimated phenotypes. Overall, this model describes the core metabolic pathways specific to *N. oceanica*, which allows for specifically tailored metabolic target predictions. With relatively few constraints we could reproduce carbon assimilation at varying light conditions and differentiate between light-acclimated phenotypes. Moreover, localization predictions have been included in the GEM to support target protein selection. Although network predictions are relatively under-constrained and can be refined further, this GEM provides a first overview of the *N. oceanica* metabolism, which can be used to further understand and improve *N. oceanica* as photosynthetic chassis for lipid production.

## Data availability statement

The model iSO1949_N.oceanica is accessible from BioModels (Malik-Sheriff et al., 2020) under identifier MODEL2503190001 and accessible on GitLub in StandardGEM format (Anton et al., 2023) on https://gitlab.com/wurssb/Modelling/i-so-1949-n-oceanica-gem where community efforts for improving the GEM can be added continuously.

