## Appendix S1 for "From Light to Lipids: Constraint-Based Metabolic Modeling of *Nannochloropsis oceanica* Under Light Acclimation Conditions"

Appendix to the article "Light to Lipids: Constraint-Based Metabolic Modeling of *Nannochloropsis oceanica* Under Light Acclimation Conditions", by van Oossanen et al. [under review]

#### A. Pipeline for protein localization predictions

In the following sections we provide supplementary information on our approach for predicting protein localizations. First, all putative coding sequences of *N. oceanica* IMET1 were scored using tools and genomic input described in the main text and Supplementary file S2A. Second, each protein was given a score per compartment, depending on compartment specific tool predictions (Supplementary file S2E-H) (Table S1.1). Third, the protein was assigned to a compartment based on the highest compartment score. A visual summary of the localization is provided in Supplementary file S2B, and Figure S1.1.

Nuclear scores were determined based on a positive prediction of predictNLS and a DeepLoc prediction containing 'Nucleus'. Positive hit to either tool increases the prediction score by 1, resulting in a maximum score of 2 for the nucleus.

Chloroplast localization scores were determined based on SignalP4.1, 5, and 6, ASAFind, ASAFind 2.0, TargetP 2.0 and DeepLoc 2.0. ASAFind was run using SignalP4.1 as input, and ASAFind 2.0 with TargetP 2.0 as input. When a positive signal peptide hit was found by any SignalP version, we considered SignalP to be positive. Also, presence of the F, W, Y, or L on the +1 position of the cleavage site was assessed, using cleavage site predictions by SignalP and TargetP. Presence of F, W, Y, or L on the cleavage sites, SignalP predictions, and TargetP predictions were used to compute chloroplast localization predictions using the pipeline provided by Levering et al. (Levering et al., 2016). Scores would be assigned based on positive chloroplast prediction with DeepLoc 2.0 (1), the Levering et al. pipeline (1), and the ASAFind score (2 for 'low confidence', 3 for 'high confidence' or 'PPC'). ASAFind was weighed more heavily compared to other predictions since this tool is specifically developed for heterokont microalgae. When ASAFind and ASAFind 2.0 provided different outcomes, the highest confidence was used, in the order of 'low confidence', 'PPC', 'high confidence'. The maximum chloroplast score is 5.

A comparison between the outputs of ASAFind and ASAFind 2.0 is provided in Table S1.2. Overall, 810 proteins were identified as plastidic by both versions, but 483 proteins were indicated as plastid by only one of the versions. Using ASAFind 2.0 outputs over those of the initial ASAFind resulted in an increase of chloroplast localization predictions, with higher certainty as observed by the increased chloroplast scores (Figure S1.2). The prediction scores were increased further when considering the highest prediction, or consensus score, of both tools.

Endoplasmic reticulum (ER) predictions are based on DeepLoc 2.0, SignalP4.1, 5.0, and 6.0, and TargetP 2.0 predictions, in addition to the ER signal peptide [K/D][D/E]EL sequence. The signal peptide, SignalP, and TargetP predictions were used to compute ER localization predictions using the pipeline provided by Levering et al. (Levering et al., 2016). The ER localization score is based on presence of the ER signal peptide (3), DeepLoc (2), and on presence of a signal peptide according to SignalP in absence of a chloroplast signal peptide (1). The maximum ER score is 6.

Mitochondria, peroxisome, cytoplasm, and cytoplasm/ER predictions were predicted based on SignalP, DeepLoc, TargetP, HECTAR, and MitoFates predictions, in addition to presence of a peroxisome signal peptide sequence [S/A/C][K/R/H][L/M] or [SSL]. We used the Levering et al. (Levering et al., 2016) pipeline to predict localizations for each of these compartment categories, where we replaced Mitoprot with MitoFates which shows increased performance (Fukasawa et al., 2015). Mitochondria scores were assigned based on DeepLoc (1) and pipeline mitochondrial predictions (2). Peroxisome scores were assigned based on DeepLoc (1), the pipeline (2), and presence of the peroxisomal signal peptide (1). Proteins that had not received any localization score were assigned to the cytosol. A score of 1 was assigned in case of a DeepLoc cytosol prediction.

*Table S1.1 Localization score distributions of the nuclear encoded proteins (n = 12442), for each compartment. Scores and localizations are assigned according to the extended pipeline developed in this study. Higher scores indicate increased predictive support by the localization tools. Proteins assigned to the cytosol/ER are included in the Cytoplasm compartment proteins with a localization score of '1'.*

| | | Compartment | | | | | |
|---|---|---|---|---|---|---|---|
| | | Chloroplast | Mito | ER | Peroxisome | Nucleus | Cytoplasm |
| Protein localization score | 0 | 10633 | 11152 | 10933 | 12204 | 8543 | 2632 |
| | 1 | 773 | 691 | 1285 | 65 | 3428 | 9811 |
| | 2 | 233 | 80 | 177 | 3 | 471 | |
| | 3 | 131 | 519 | 41 | 136 | | |
| | 4 | 215 | | 3 | 34 | | |
| | 5 | 457 | | 1 | | | |
| | 6 | | | 2 | | | |

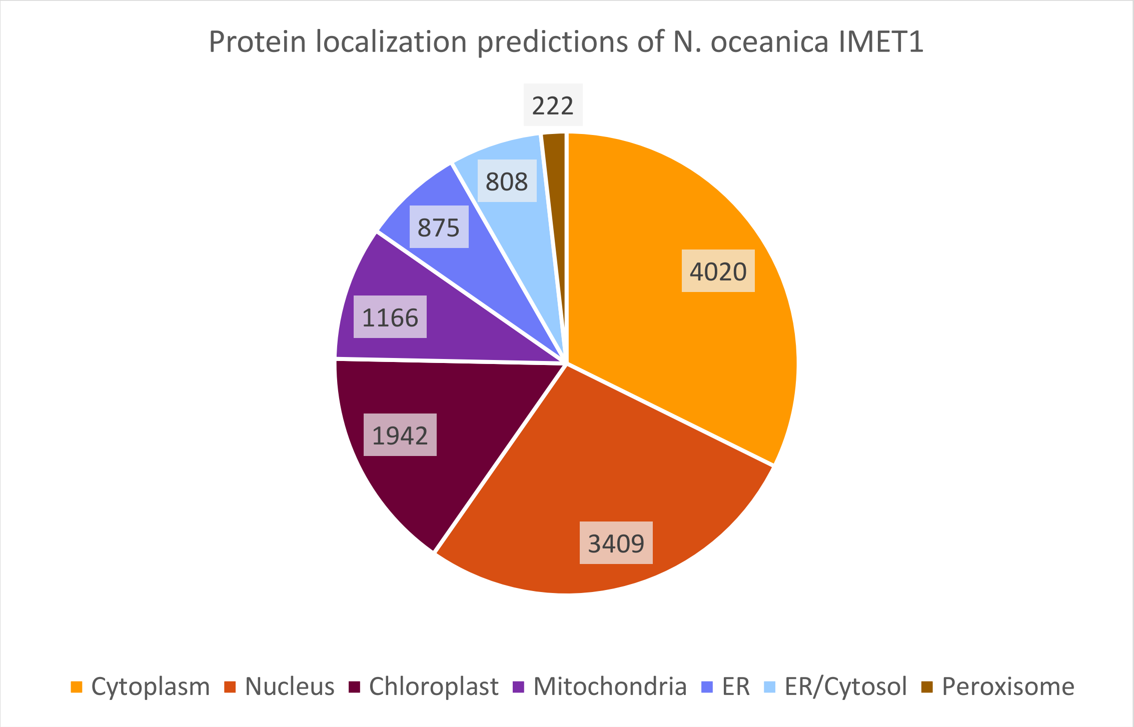


*Figure S1.1 Number of assigned proteins per compartment of the iSO1949_N.oceanica model,, using the annotated protein encoding sequences of* N. oceanica *IMET1, version 2 (Gong et al., 2020)*

*Table S1.2 Comparison ASAFind 1.0 with SignalP4.1 as input, and ASAFind 2.0 with TargetP2.0 as input, on the proteome* N. oceanica *IMET1 v2*

| ASAFind outputs (>), ASAFind 2.0 outputs (below) | Plastid, high confidence | Plastid, low confidence | Not plastid, SignalP positive | Not plastid, SignalP negative |
|---|---|---|---|---|
| Plastid, high confidence | 575 | 20 | 10 | 133 |
| Plastid, low confidence | 27 | 183 | 22 | 97 |
| PPC | 5 | 0 | 85 | 19 |
| Not plastid, not PPC, signal peptide identified | 14 | 21 | 611 | 122 |
| TargetP-2.0: mTP | 11 | 10 | 22 | 538 |
| TargetP-2.0: noTP | 29 | 32 | 76 | 9780 |

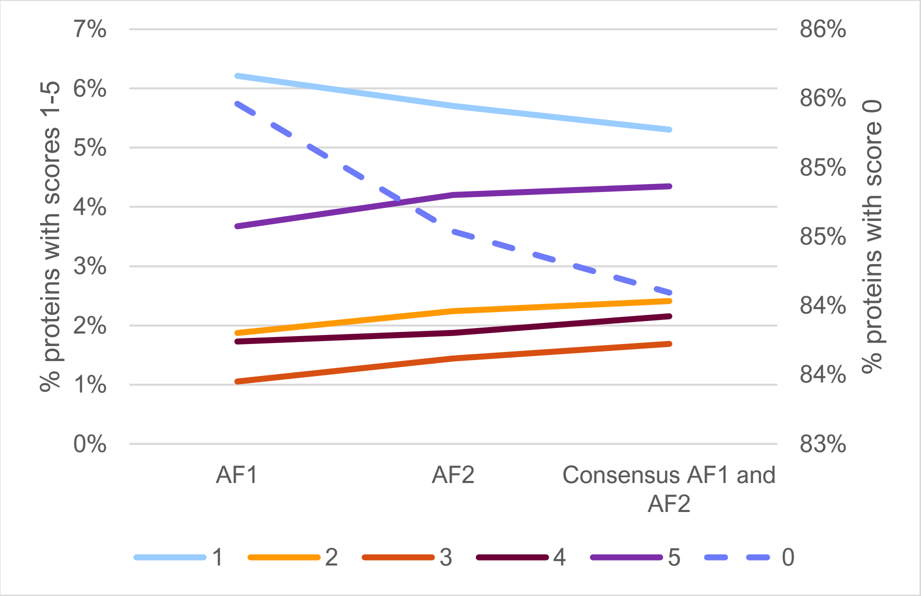


*Figure S1.2 Protein percentage with respective chloroplast score, according to ASAFind 1, 2, or taking the highest score of both*

### B. GEM draft construction

#### B.1 Modifications to AuReMe inputs: scaffold GEMs and protein sets

##### B.1.1 iLB1027_lipid, Phaeodactylum tricornutum

The nuclear encoded protein sequences were obtained from EnsemblProtists database, based on the genome assembly ASM15095v2 (https://ftp.ensemblgenomes.ebi.ac.uk/pub/protists/release-57/fasta/phaeodactylum_tricornutum/pep/ ). Plastidic and mitochondrial protein sequences were obtained from (Oudot-Le Secq et al., 2007; Secq and Green, 2011). The GEM file was obtained from the BiGG database (http://bigg.ucsd.edu/models/iLB1027_lipid).

Several locus tags were inconsistent between the protein sequence files and the GEM file. Locus tags in the GEM file were updated by changing "PHATRDRAFT_[x]" to "Phatr3_J[x]". A single gene was present in the GEM but absent in the protein sequences, AEZ63317. The AEZ63317.1 protein sequence was retrieved from NCBI and added to the protein list. The GEM file was edited one last time to enable AuReMe to read the GPR by including it in the reaction notes as "GENE_ASSOCIATION: [gpr]."

After orthology-based draft construction, edits made to iLB1025 to form iLB1034 were included in the draft GEM.

##### B.1.2 iRJ1321, Nannochloropsis gaditana

The nuclear, mitochondrial and plastidic encoded protein sequences were obtained from the Genbank (GCA_000569095.1). The GEM file was obtained from the article and preprocessed for AuReMe by removing reaction unassociated genes.

##### B.1.3 iNS934, Nannochloropsis salina

Both the protein sequences and GEM file were obtained from the article and preprocessed for AuReMe through the following edits:

- GPR " [..] and and 537_psba_cp [..]" was changed to "[..] and 537_psba_cp [..]" in the SBML file
- All "evm.m1[x]" GPR (24 genes) were changed to "evm.model[x]" in the SBML file
- "cds.NSV[x]|m.[x]" was changed to "NSV[x].m.[x]" in the protein sequence file
- Protein sequence "NSV4002001.m.17653" was lacking, and could not be retrieved within this study.
- Parentheses in metabolite and reaction IDs were replaced by underscores.

##### B.1.4 iCre1355, Chlamydomonas reinhardtii

Protein sequences were obtained from JGI, *Chlamydomonas reinhardtii* v5.5 (https://phytozome-next.jgi.doe.gov/info/Creinhardtii_v5_6), the SBML file from the article ("iCre1355_auto.xml").

#### B.2 Conversion of draft GEM padmet to SBML

In order to convert the resulting iNoce draft.padmet to an SBML file, several metabolite ID edits were made to the padmet file:

- M_4__methyl/cholestadienol_c -> M_4__methyl_cholestadienol_c
- M_pseudouridine_5_p_c -> M_pseudouridine5p_c
- M_adenosylcobalamin_5_p_h -> M_adenosylcobalamin5p_h
- M_2_3_dihydrodipicolinate_c -> M_23dihydrodipicolinate_c

In addition, all underscores in all IDs were replaced by a placeholder letter string, and reverted to underscores after sbml construction.

### C. Metabolite mapping

#### C.1 BiGG database mapping

Metabolites from the orthology-based draft reconstructed draft GEM were mapped using firstly the BiGG database. 3459 out of 4718 metabolite IDs from the draft GEM were found in either the BiGG IDs or in the mapping IDs associated with BiGG metabolites, with the following distribution:

| Number of metabolite ID matches with [annotation] present in the BiGG database: | | | | | | |
|---|---|---|---|---|---|---|
| **No. metabolites of iNoce draft** | **Universal BiGG ID** | **Out-of-use BiGG ID** | **Metacyc mapping** | **Seed mapping** | **KEGG mapping** | |
| 211 | x | | | | | BiGG ID present |
| 28 | | x | | | | old BiGG ID |
| 3 | x | x | x | | | BiGG ID |
| 3038 | x | x | | | | BiGG ID |
| 148 | | | x | | | Mapped to BiGG from Biocyc |
| 0 | | | | x | | Mapped to BiGG from ModelSEED |
| 31 | | | | | x | Mapped to BiGG from KEGG |
| 1259 | | | | | | Mapped manually or with MetaNetX |

#### C.2 BioCyc Database mapping

Additionally, mapping of 1915 metabolites could be supplemented with Biocyc and MetaNetX automatic mapping.

Unique metabolite IDs (without compartment tag) were submitted several times to BioCyc for each of the following settings:

- IDs not defined to a DB
- IDs defined as BiGG IDs
- IDs defined as KEGG IDs
- IDs defined as Seed IDs
- IDs defined as Biocyc IDs

Results from ‘IDs defined as Biocyc’ and ‘IDs not defined to a DB’ are identical, no results were retrieved when draft metabolite IDs were defined as ‘Seed IDs’.

### D. Orthology-based GEM construction

#### D.1 FAIR curation and lipid labelling

Extensive mapping of the metabolite IDs and curation of reactions resulted in removal of 812 metabolites and 1292 reactions. In this way, metabolites appearing multiple times with different identifiers or reactions with conflicting substrate or product usage were largely eliminated. Additionally, 970 metabolites and 1663 reactions were removed to simplify lipid biosynthesis and limit lipid metabolic reactions to the species with single acyl-chain types detected in our study. Removed reactions and metabolites involving lipids with mixed acyl-chain types are listed separately in Supplementary table S4 and can be added again to the model in case more detailed lipid composition data becomes available. Consistent mapping of the metabolite and reaction IDs further increased the number of overlapping reactions originating from multiple scaffolds by 308. As a result, out of a total of 3485 reactions in the curated iSO1949_N.oceanica GEM, 1271 reactions are supported by orthology with multiple scaffold organisms.

#### D.2 Contribution of scaffold GEMs to *N. oceanica* IMET1 GEM based on orthology

We compared the relative contribution of each scaffold GEM to the iSO1949_N.oceanica draft and final GEM construction, based on the number of orthologous reactions per scaffold GEM (Figure 3, Supplementary table S4.2 ). All scaffold GEMs contributed a similar number of reactions to the iSO1949_N.oceanica draft (1590-2215), despite varying strongly in degrees of orthology between the scaffold GEMs associated genes and the *N. oceanica* genome (53%-99%). The iLB1027 GEM provided the most orthologous reactions to the *N. oceanica* GEM draft (2215) during orthology-based reconstruction, despite having the lowest orthology rate out of all 4 scaffold GEMs (covering 53% of iLB1027 reactions with GPR). On the other hand, almost complete orthology was reached by the *Microchloropsis* GEMs; respectively 96% and 99% of the iRJ1321 and iNS934 reactions were orthologous to *N. oceanica* genome.

During curation of the draft GEM, reaction contributions per scaffold were reduced to between 1231 and 1565 reactions. Interestingly, curation of the iSO1949_N.oceanica strongly reduced the contribution of iLB1027 to the GEM from 2215 to 1391 reactions, mainly due to simplification of the lipid biosynthesis pathways. On the other hand, iRJ1321 of *M. gaditana* contributed most reactions after curation and was reduced least owing mostly to a similar level of simplicity in organelles and lipid biosynthesis. Before curation 1594 reactions of iRJ1321 were included, after curation 1565, which is 48% of the total reactions in iSO1949_N.oceanica.

Manual curation tailored the list of reactions to *N. oceanica*, and revealed increased similarity between the transferred reactions of the scaffold GEMs. Initially, little overlap was seen between orthologous reaction IDs from each scaffold, except for iNS934 and iCre1355 which overlapped with more than 50% of their reactions (938 out of 1719 and 1590 reactions resp.) (Figure 3, left). Manual curation increased connectivity between the reactions from the different scaffolds, especially between the *M. gaditana* iRJ1321 and the other scaffolds GEMs where 800 out of 1565 reactions showed overlap after curation compared to no overlap before curation (Figure 3, right). The number of shared reactions between the other scaffold GEMs also increased from 217-965 before curation, to 654-1066 after curation despite the removal of irrelevant compartments and pathways. Still, differences in contribution can be seen strongly for *M. gaditana* and *P. tricornutum* which provided 765 and 737 unique reactions respectively. The unique reactions contributed by *M. gaditana* and *P. tricornutum* GEMs are mostly related to the lipid metabolism, 302 and 342 unique reactions respectively, despite the strong simplification and reduction of the lipid biosynthesis pathway provided by iLB1027 through orthology. The *P. tricornutum* GEM iLB1027 is thoroughly curated on the lipid metabolism, containing relatively higher numbers of lipid metabolic reactions (1856 compared to 788, 536, and 616 of iNS934, iRJ1321, and iCre1355 resp.).

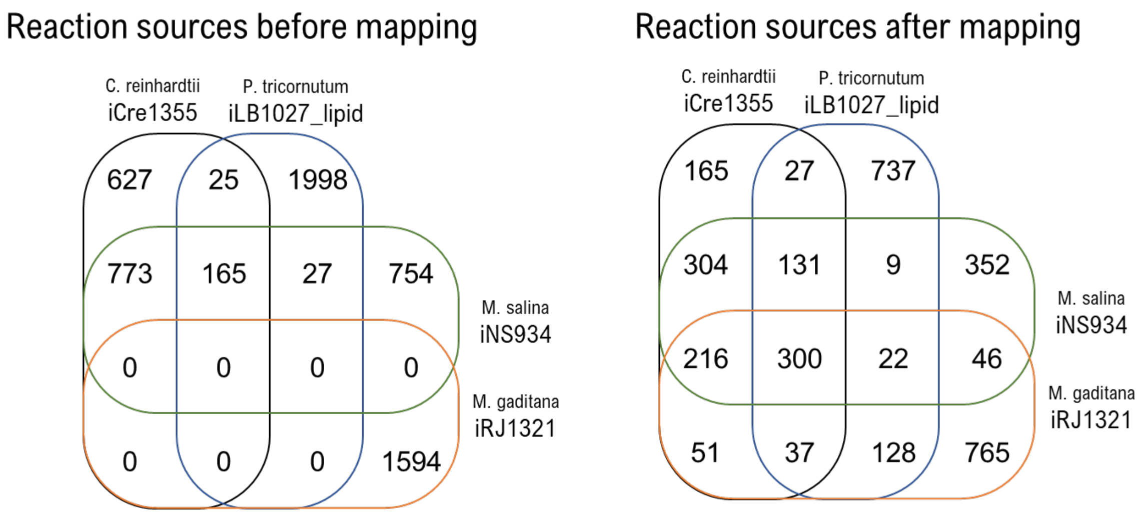


*Figure 1.3 Scaffold contribution to draft GEM after automatic GEM construction (left) and to the final iSO1949_N.oceanica GEM after manual curation (right, see Figure 3 of the main text), based on reaction IDs.*

### E. GEM reaction localization versus predicted locations

#### E.1 Localization predictions relative to GEM annotation

Often multiple genes are assigned to certain reactions. We assessed whether reaction compartments matched the predicted localization of their gene-associated protein products, also called Gene-Protein-Reaction (GPR). For each GEM compartment, we evaluated the predicted location of the proteins associated to the reactions (section 1A). A substantial match between predicted GPR location and reaction compartments was seen for the cytosol, thylakoid, and mitochondria. However, GEM reactions in the ER showed discrepancy, since only 40% of the reactions were related to an ER predicted protein product, as compared to 61% with at least one cytosol predicted protein in their GPR. A significant factor contributing to these mismatched protein localizations is the large number of lipid biosynthesis reactions currently located in the ER GEM compartment. Lipid biosynthesis is suggested to occur partly in compartments between the chloroplast membranes, from which the outer membrane is merged with the ER. Localization prediction of proteins to these compartments is currently not possible, although efforts are being made to predict localization to the periplastidic compartment (PPC) between the second and third chloroplast membrane (Gruber et al., 2023). Consequently, to maintain continuity between lipid biosynthesis and degradation pathways, most algal GEMs assign lipid metabolic reactions to the cytosol or ER. A similar trend to the ER reactions is observed for chloroplast reactions, with 30% of the associated proteins predicted to be ER-localized and 47% cytoplasm-localized. Conversely, peroxisomal GEM reactions were predominantly associated with peroxisomal-predicted proteins (66%), but almost equally with ER-predicted proteins (67%) as well.

Despite the apparent discrepancy between reactions and predicted GPR localizations for certain compartments, between 98-100% of the reactions with GPR had at least one gene product matching in location (section 1A). This is an improvement compared to protein predictions using the Levering pipeline (Levering et al., 2016), which results in reduced matches in the chloroplast and thylakoid thereby reaching 88-94% overlap for these compartments.

*Table S3 Match between reaction locations in GEM and the predicted localization of their GPR. Percentages indicate the GPR ratio of reactions in compartment (row) predicted to be in a certain compartment (column). For example, from the 2548 cytosolic reactions, 73% had at least one gene in the GPR predicted to be in the cytoplasm. Compartment abbreviations: c – cytosol, h – chloroplast, u – thylakoid, r – endoplasmic reticulum, m-mitochondria, x – peroxisome, n – nucleus.*

| | | no. rxns | Predicted localization of reaction GPR | | | | | |
|---|---|---|---|---|---|---|---|---|
| | | | **Cytoplasm** | **Plastid** | **ER** | **Mitochondria** | **Peroxisome** | **Nucleus** |
| **Reaction compartment** | **c** | **2545** | 73% | 30% | 30% | 18% | 8% | 25% |
| | **h** | **1211** | 47% | 54% | 30% | 24% | 14% | 14% |
| | **u** | **24** | 25% | 75% | 0% | 8% | 0% | 0% |
| | **r** | **162** | 61% | 14% | 40% | 16% | 2% | 29% |
| | **m** | **587** | 42% | 25% | 23% | 75% | 15% | 6% |
| | **x** | **121** | 45% | 2% | 67% | 13% | 66% | 3% |
| | **n** | **15** | 40% | 40% | 0% | 13% | 0% | 53% |

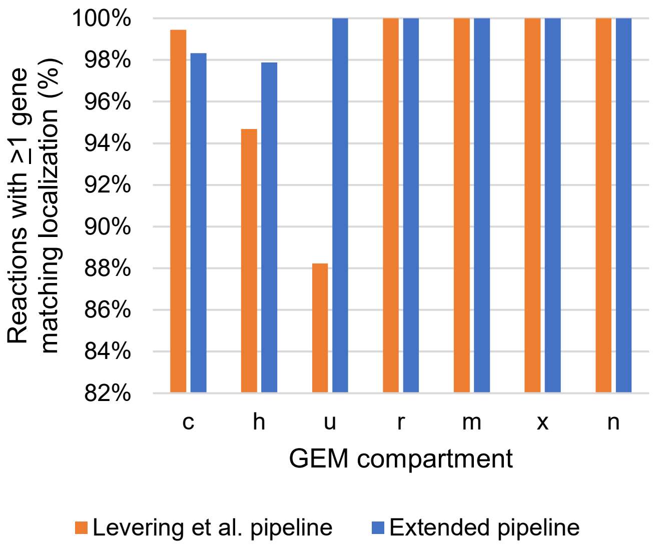


*Figure S4 Ratio of reactions per compartment, with at least 1 associated gene matching in localization prediction with the respective model compartment. Localization was predicted of the N. oceanica annotated proteins according to the approach of Levering et al. pipeline (Levering et al., 2016). Compartment abbreviations: c – cytosol, h – chloroplast, u – thylakoid, r – endoplasmic reticulum, m- mitochondria, x – peroxisome, n – nucleus.*
